# Metabolic Reprogramming of Pathogenic CD4^+^ T Helper Cells Attenuates Inflammatory Bowel Disease Pathogenesis

**DOI:** 10.1101/2025.04.15.649047

**Authors:** Shravan K. Mishra, Leena M. Abdelrahman, Heidi M. Davidson, Hyun Se Kim Lee, Lucía Valenzuela-Pérez, Omar M. Hassan, William A. Faubion, Petra Hirsova, Adebowale O. Bamidele

**Author notes:** Correspondence Address correspondence to: Adebowale Bamidele, Ph.D., Division of Gastroenterology and Hepatology, Mayo Clinic, 200 First Street SW, Rochester, Minnesota 55905, USA.

## Abstract

**BACKGROUND & AIMS:** CD4^+^ T helper 1 (Th1) cells are involved in human inflammatory bowel disease (IBD) pathogenesis; however, mechanisms governing the persistent inflammatory function of these cells are unclear, leading us to examine how metabolism governs Th1 cell-induced IBD.

**METHODS:** Th1 cells supplemented with methyl pyruvate (MePyr) were analyzed to define how enforced mitochondrial pyruvate metabolism and subsequent glycogen synthase kinase 3β (GSK3β) deactivation reprogram cellular state. Re-analysis of the inflamed ileal single-cell RNA sequencing dataset from Crohn’s disease patients was performed to assess non-Treg CD4^+^ T cell metabolic gene signature. We assessed the capacity of a repurposed GSK3β inhibitor to restrain pathogenic CD4^+^ T cell-driven murine colitis.

**RESULTS:** Effector Th1 cells exhibit a distinct metabolic program exemplified by glucose-driven glycolysis but low mitochondrial respiration. MePyr deactivates GSK3β, glycolysis, and histone H3 acetylation on cytokine promoter region, resulting in reduced interferon-γ (IFN-γ) and tumor necrosis factor-α (TNF-α) expression in Th1 cells with concomitant gain of regulatory T cell-like program. GSK3β inhibition with LY2090314 mirrored the anti-inflammatory effect of MePyr in a manner reversible by acetate supplementation, implying that GSK3β potentially sustains glycolysis-derived acetyl-coenzyme A needed for histone acetylation and Th1 cell inflammatory response. Interleukin-21 exacerbates Th1 cell inflammatory response by maintaining a GSK3β-driven glycolytic program. The Th1 cell metabolic gene signature downregulated by MePyr or GSK3β inhibition *in vitro* is enriched in refractory Crohn’s disease patients. GSK3β inhibition with LY2090314 retrains T cell-induced colitis in mice.

**CONCLUSIONS:** MePyr impairs GSK3β-mediated glycolysis and Th1 cell immune response. GSK3β inhibition may mitigate Th1 cell-induced human IBD.

**Graphical abstract:** 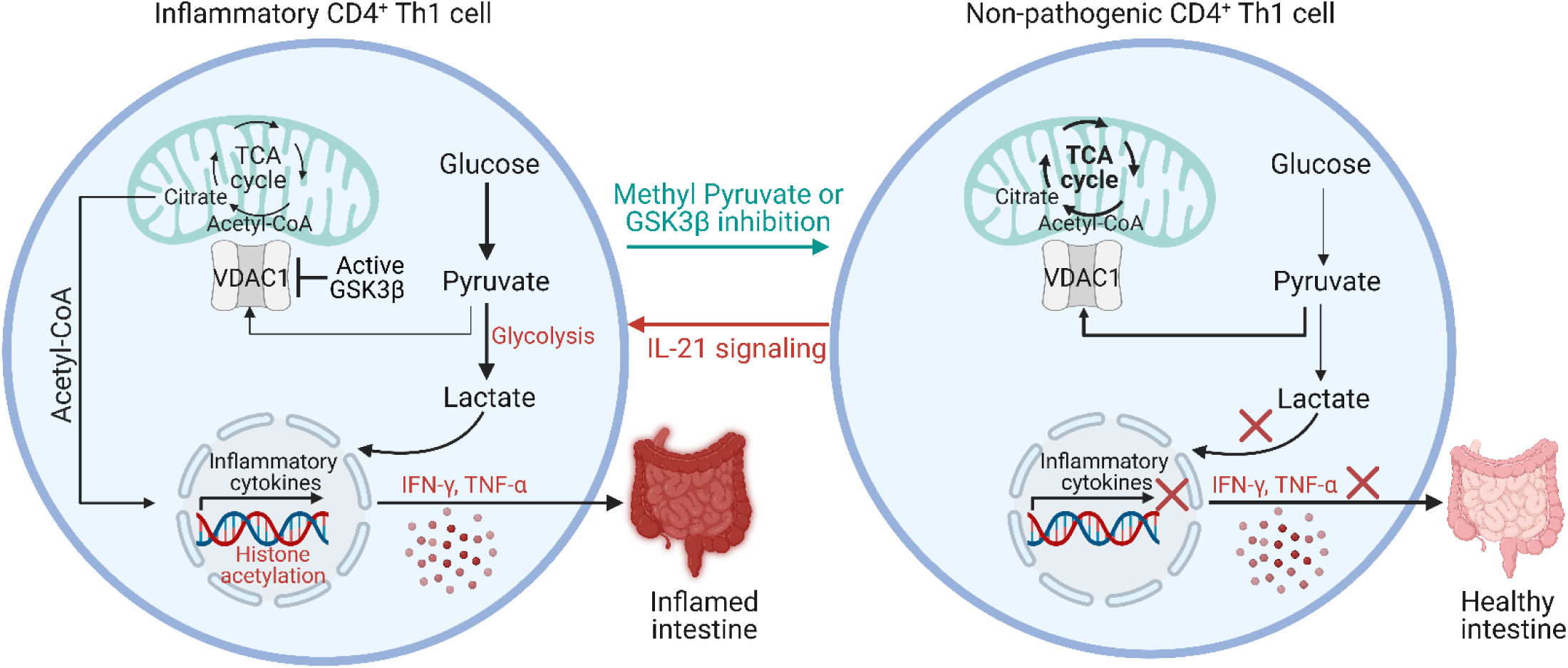

## Introduction

Cellular metabolism is a highly dynamic process that supports many aspects of the cell’s activities.^1, 2^. Metabolism is orchestrated by metabolic enzymes organized into pathways specialized for processing and producing distinct metabolites in a compartmentalized fashion in cellular organelles. Different pathways share common metabolites and critical bifurcation points in complex biochemical networks. A well-appreciated example of a metabolic bifurcation point is the metabolism of glucose-derived pyruvate in the glycolysis pathway or by the mitochondrial tricarboxylic acid (TCA) cycle linked to citrate to acetyl-Coenzyme A (acetyl-CoA) shuttle system or oxidative phosphorylation (OXPHOS) and ATP production in the mitochondria^1, 2^. Here, we explore the metabolic underpinnings of CD4^+^ T cell inflammatory function and the therapeutic consequence of limiting metabolic flexibility by supplementing Th1 and Th17 cells with a mitochondria membrane-permeable form of pyruvate.

Effector CD4^+^ T cells are integral to the adaptive immune response, differentiating into T helper type 1 (Th1), Th2, and Th17 cell subsets tuned to respond to a wide range of pathogens and environmental insults^3^. Th1 cells produce the signature cytokine interferon (IFN)-γ that functions to eradicate intracellular pathogens^4, 5^. Th17 cells typically secrete interleukin-17 (IL-17A/F) cytokines at mucosal surfaces under homeostatic conditions where they provide protection from pathogenic bacteria and fungi^6–8^. Th1 and Th17 cell responses are tightly controlled to prevent host tissue damage following pathogen elimination. However, under conditions that favor chronic inflammatory processes, Th1 and Th17 cells can adopt pro-inflammatory programs, typified by IFN-γ and tumor necrosis factor (TNF)-α production that promotes immune-mediated diseases, such as inflammatory bowel disease (IBD; mainly Crohn’s disease and ulcerative colitis)^9–13^. Therefore, understanding how damaging microenvironmental cues sustain cellular metabolic events associated with persistent immune response by effector Th1 and Th17 cells may lead to the development of efficacious therapies.

Metabolite tracing studies have revealed that activated T cells decrease the rate of glucose-derived pyruvate entry into mitochondria in favor of aerobic glycolysis (conversion of pyruvate to lactate)^14, 15^. To meet cellular demands, T cell activation is accompanied by OXPHOS to glycolysis metabolic shift that provides key intermediate metabolites required for the acquisition of effector function^16–18^. Despite the decreased utilization of glucose-derived pyruvate for mitochondrial metabolism by effector CD4^+^ T cells, aerobic glycolysis has been shown to contribute to IFN-γ production by increasing cytosolic acetyl-CoA pools via the export of glucose-derived mitochondrial citrate to the cytoplasm^19, 20^. Despite these findings, our understanding of the molecular switches that control how CD4^+^ T cells acquire effector function is incomplete. In this study, we aimed to identify metabolic checkpoints required for Th1 and Th17 cell effector function by enforcing mitochondrial pyruvate metabolism in these cells.

Here, we developed models of human effector CD4^+^ T cells that allowed us to interrogate pathways driving inflammatory cytokine expression. We found that enforced mitochondrial pyruvate metabolism through supplementation of effector Th1 cells with membrane-permeable methyl pyruvate attenuated glycolysis, leading to attenuated Th1 cell inflammatory program and concomitant acquisition of a regulatory T cell-like phenotype. Mechanistically, MePyr suppressed glycogen synthase kinase 3β (GSK3β) enzymatic activity, which expectedly activated voltage-dependent anion channel-1 (VDAC1), leading to a reduction in glycolysis, intracellular acetyl-CoA, histone H3 acetylation, and inflammatory cytokine expression. Transcriptional analysis of the single-cell RNA sequencing dataset from refractory Crohn’s disease patients revealed enrichment of glycolytic genes in the inflamed ileal non-Treg CD4^+^ T cells. In a disease setting, GSK3β pharmacologic inhibition restrained pathogenic CD4^+^ T cell-induced colitis in mice.

## Materials and Methods

### Healthy Blood Donors

After written informed consent, we obtained blood samples from healthy male donors (19-60 years old) as apheresis cones. Buffy coats were obtained from apheresis cones (Mayo Clinic Blood Donor Center).

### Animals

C57BL/6J *Rag1*^-/-^ mice were purchased from the Jackson Laboratory and kept in conventional housing in the Mayo Clinic animal facility. C57BL/6NJ WT and *Il21r*^-/-^ (*Il21r^tm^*^1k^*^opf^*) mice were purchased from the Jackson Laboratory. All mice were housed in the Mayo Clinic animal facility. Mice used in experiments were males of 6– 9 weeks of age. We performed all animal work in accordance with reviewed/approved protocol by the Mayo Clinic Institutional Animal Care and Use Committee.

### CD4^+^ T cell Isolation from Human PBMCs and Mouse Spleen, Cell Culture and T Cell Differentiation

Naive CD4^+^ T cells were isolated from male mouse spleen (CD62L^+^) or human PBMCs (CD45RA^+^) using the corresponding CD4^+^ T cell isolation kit according to the manufacturer’s instructions (Miltenyi Biotec). All primary human and mouse CD4^+^ T cells were cultured under standard conditions with cRPMI 1640 (RPMI 1640 + L-Glutamine [Gibco] supplemented with 10% FCS, 10 mmol/L HEPES pH 7.4, 1 mmol/L sodium pyruvate, 2 mmol/L L-glutamine, 1% non-essential amino acids, penicillin, streptomycin, and 50 µmol/L β-mercaptoethanol). Human CD4^+^ T cell subsets were polarized from naïve cells in the presence of anti-human CD3 (2 µg/ml) and CD28 (2 µg/ml), and human IL-12 (20 ng/ml) for Th1 cells, human IL-4 (10 ng/ml), IL-2 (100 U/ml), and anti-human IFN-γ (10 µg/ml) for Th2 cells, human TGF-β1 (0.5 ng/ml), IL-1β (20 ng/ml), IL-6 (50 ng/ml), IL-21 (100 ng/ml), IL-23 (20 ng/ml), and anti-human IL-4 (2.5 µg/ml) and IFN-γ (5 µg/ml) antibodies for Th17 cells, and human TGF-β1 (5 ng/ml) and IL-2 (100 U/ml) for iTregs for 5 days with fresh media and cytokines added on day 2 and day 4. On day 5, differentiated T cells were harvested, resuspended in fresh media only and plated for 2-days in the presence of anti-human CD3 (2 µg/ml) and CD28 (2 µg/ml) antibodies. Differentiated (day 5) and developed (day 7-8) human CD4^+^ T subsets were treated with indicated inhibitors or metabolites in regular cRPMI medium for the indicated time points. Murine Th1 cells were obtained by culturing naive CD4^+^ T cells for 4 days in the presence of anti-mouse CD3 (2 µg/ml), anti-mouse CD28 (2 µg/ml) and recombinant murine IL-12 (25 ng/ml) and human IL-2 (100 U/ml), and anti-mouse IL-4 (2.5 µg/ml) and IFN-γ (5 µg/ml) antibodies, while murine Th17 cells were obtained by culturing naïve cells in the presence of recombinant murine IL-6 (25 µg/ml), murine IL-23 (25 ng/ml), human TGF-β1 (1 ng/ml), anti-mouse IL-4 (2.5 µg/ml) and IFN-γ (5 µg/ml) antibodies.

### Colitis Induction and Prevention in Rag1 Knockout (T and B-cell deficient) Mice

Male C57BL/6J mice and C57BL/6J *Rag1*^-/-^ mice (Jackson Laboratory) were placed in conventional housing at the Mayo Clinic animal facility. For CD4^+^ CD45RB^high^ T cell-induced colitis induction, CD4^+^ T cells were isolated from splenocytes of normal wild-type (WT) C57BL/6J mice. CD4^+^ T cells were co-stained with CD45RB and CD25 fluorescently conjugated antibodies and sorted for CD4^+^CD45RB^high^ T cells and CD4^+^CD25^++^ (Tregs) via flow cytometry. 500,000 CD4^+^ CD45RB^high^ T cells were injected into *Rag1*^-/-^ mice intraperitoneally (i.p.) along with vehicle control (4% DMSO 45% PEG-300), GSK3β inhibitor LY2090314 (Selleckchem: 10 mg/kg) in 45% PEG-300, or 500, 000 Treg cells on day 0 to treat or prevent disease. The weights of experimental mice were monitored for 0-12 weeks while being fed a non-irradiated chow diet. Mice used in experiments were males of 6–9 weeks of age. All animal work was done in accordance with and reviewed/approved by the Mayo Clinic Institutional Animal Care and Use Committee.

### Colitis Assessment and Histopathology

Mice were weighed every week and their colon lengths were determined during necropsy. The degree of colitis was quantified using three outcome variables: weight loss, colon histology, and disease activity index. Mouse Colon Histology Index (MCHI) assesses eight parameters, including goblet cell loss, crypt density, crypt hyperplasia, muscle thickening, crypt abscess, ulceration, the extent of inflammatory infiltrate, and semiquantitative assessment of submucosal inflammation. The Disease Activity Index (DAI) is an established clinical index of colitis severity encompassing clinical signs of colitis (wasting and hunching of the recipient mouse and the physical characteristics of stool) and an ordinal scale of colonic involvement (thickness and erythema). The dissected colon was fixed in 10% neutral buffered formalin for 24 hours and embedded in paraffin. Tissue sections (4 µm) were cut using a microtome and positioned on glass slides. Hematoxylin and eosin (H&E) staining were performed according to standard techniques. H&E slides were reviewed by a blinded gastrointestinal pathologist.

### Serum Cytokine Analysis

Cytokine levels were determined in supernatants (50 µl of serum) using the BD cytometric bead array mouse Th1/ Th2/Th17 kit (BD Bioscience) according to the manufacturer’s instructions and analyzed using FCAP Array version 3 software (Soft Flow Hungary Ltd., Pécs, Hungary).

### Proximity Ligation Assay (PLA) and Confocal Microscopy of Fixed Cells

As previously described ^21^, cells were harvested and plated on 8-well Lab-Tek chamber slides (Thermo Fisher Scientific, Waltham, MA) coated with fibronectin (1 mg/ml) (Corning, Corning, NY) for 3 hours to allow cell attachment and then stimulated or treated as indicated. Cells were stained with organelle tracking dyes (200-500 nM range) and then fixed with 4% paraformaldehyde, permeabilized with 0.15% Triton X-100 (Invitrogen, Carlsbad, CA), and washed with PBS. Cells were blocked for 1 h with 5% bovine serum albumin containing 0.1% glycine and incubated with anti-VDAC1 and anti-IP3R1 antibodies to biochemically measure mitochondria-ER junctions of < 30 nm distance. Mitochondria-ER junctions were visualized by Duolink in situ fluorescence PLA probes and detection reagents in accordance with the manufacturer’s instructions (Sigma, St. Louis, MO). Cells were mounted with Duolink 40,6-diamidino-2-phenylindole–containing mounting (DAPI) medium (Sigma-Aldrich). Images of cells were captured by using a C-Apochromat 63_ objective/1.20 W korrM27 of a fluorescent confocal microscope (LSM780 AxioObserver; Carl Zeiss, Jena, Germany). Images were processed using the Zen lite 2012 software (Carl Zeiss). Quantification of detected PLA signals or dots was measured using ImageJ (National Institute of Health, Bethesda, MD).

### Extracellular Flux Analysis using Agilent Seahorse XF Analyzer

XFe-96 Extracellular Flux Analyzer (Seahorse Bioscience, Agilent Technologies) was used to define OCRs and ECARs. 250,000-300,000 cells were seeded on XF96 Cell Culture Microplates before perturbation experiments. Mitochondrial perturbation experiments with cells were conducted in XF media containing glucose (25 mM or as otherwise indicated), pyruvate (2 mM) and L-glutamine (0.5 mM) by the sequential addition of 1 μM oligomycin (Sigma-Aldrich), 1 μM FCCP (carbonyl cyanide 4-(trifluoromethoxy) phenylhydrazone; Sigma-Aldrich) and 0.5 μM rotenone/antimycin A (Sigma-Aldrich). Glycolysis stress tests with cells were conducted in XF media by the sequential addition of 10 mM glucose (Sigma-Aldrich), 2 μM oligomycin (Sigma-Aldrich), and 50 mM 2-DG (Sigma-Aldrich). Changes in OCR and ECAR in cells were measured using the Seahorse XF analyzer as indicated.

### Immunoblotting Analysis

Cells were harvested, washed with cold PBS, and lysed with Radioimmunoprecipitation (RIPA) lysis buffer containing 50 mmol/L Tris-HCl, pH 7.4, 1% NP-40, 150 mmol/L NaCl, 2 mmol/L EDTA, and protease and phosphatase inhibitors. Cell lysates were boiled in SDS sample buffer for 10 mins at 95-100° C, resolved on SDS-PAGE gel, and transferred to PVDF membranes. Membranes were blocked for 1 h in milk or BSA and then incubated with corresponding primary antibodies at 4°C overnight. After washing, membranes were incubated with secondary antibodies in TBS + Tween 20 + 5% nonfat dry milk for 1 h. ECL western blotting chemiluminescent substrates were added for 1 min and membranes were imaged on Bio-Rad imaging software. Membranes were stripped and re-probed with antibodies.

### Flow Cytometry

Cells were treated as indicated with or without brefeldin A (BioLegend), fixed with True-Nuclear Transcription Factor Buffer Fix solution (BioLegend), permeabilized with Perm/Wash buffer (BioLegend), blocked for 10 minutes and then stained with relevant fluorochrome-conjugated primary antibodies. Cells were washed twice with Perm/Wash buffer and subsequently subjected to flow cytometry for fluorescence-activated cell sorting. Cells were electronically gated on live cells for analysis.

### Gene Expression Analysis using the NanoString nCounter System

The gene expression profile of human Th1 cells was assessed using the NanoString nCounter® system (NanoString Technologies). We employed the Metabolic and Autoimmune Profiling Panels on human effector Th1 cells supplemented with MePyr 5 mM or treated with LY2090314 for 24 h. The Metabolic Profiling Panel consisted of 786 genes across 34 annotated pathways, while the Autoimmune Profiling Panel included 770 genes across 51 annotated pathways. Total RNA was isolated from human effector Th1 cells using the Zymo Research Direct-zol™ RNA MiniPrep Kit (Zymo Research) according to the manufacturer’s instructions. The quantity and quality of the extracted RNA were assessed using a NanoDrop spectrophotometer (Thermo Fisher Scientific). For the NanoString nCounter® assay, 100 ng of total RNA was hybridized at 65°C for 18 h in a thermal cycler with a custom-designed codeset specific to the target genes of interest, including housekeeping genes for normalization according to the manufacturer’s instructions. Raw data were normalized to internal positive controls and housekeeping genes to account for variability in RNA input and hybridization efficiency. Data analysis was performed using the nSolver™ Analysis Software (NanoString Technologies). Differential gene expression analysis was conducted to identify significant changes between experimental groups.

### Gene Ontology and Pathway Enrichment Analysis

After gene expression analysis was performed using NanoString Metabolic and Autoimmune Panels. Differentially expressed genes (DEGS) with a threshold of p-value <0.05 were subjected to the pathway and functional enrichment analysis using ShinyGO 0.82 (http://bioinformatics.sdstate.edu/go/), a web-based graphical tool for gene set enrichment analysis. Briefly, a list of differentially expressed genes (DEGS) was submitted to ShinyGO, using gene symbols as input. The species selected for analysis is Homo Sapiens. Enrichment analysis was conducted for Gene Ontology (GO) terms, including biological process (BP) and for pathway databases including KEGG, with a minimum pathway size of 4.

### Real-time (RT) Quantitative PCR

Total RNA was isolated with RNeasy Plus Mini Kit (Qiagen) and was reverse transcribed using the iScript cDNA synthesis kit (Bio-Rad). Quantification of gene expression was performed by real-time polymerase chain reaction using SYBR green fluorescence on a LightCycler 480 instrument (Roche). Specific primers are listed in Supplementary Table 2 in Supplementary Methods. Target gene expression was calculated using the ΔΔCt method. Expression was normalized to 18S expression levels, which were stable across all experimental groups.

### Chromatin Immunoprecipitation-quantitative PCR (ChIP-qPCR)

Chromatin Immunoprecipitation was performed as per the manufacturer’s instructions using Magna ChIP HiSens Kit (Millipore). Briefly, cells were fixed in 37% formaldehyde in phosphate-buffered saline (PBS) for 10 min and quenched with glycine for 5 min at room temperature. Cells were harvested and washed twice with PBS containing protease inhibitor cocktail. Cells were pelleted down by spinning at 800 x g at 4°C for 5 minutes, resuspended in nuclei isolation buffer containing protease inhibitor cocktails, incubated on ice for 15 minutes, and vortexed at high speed for 10 seconds every 5 minutes to enhance nuclei release. The cell suspension was spun at 800 x g at 4°C for 5 minutes and resuspended in sonication (SCW) buffer containing protease inhibitor cocktails for shearing chromatin (cross-linked Proteins/DNA) by subjecting to 8 cycles of sonication (30 s ON/OFF) at high amplitude using a Bioraptor Pico (Diagenode). Following sonication, the chromatin extract was precleared with Magna ChIP protein A/G Magnetic Beads in the sonication buffer. For each chromatin immunoprecipitation, ∼190 µL of cold SCW buffer containing Protease Inhibitor Cocktail III, 10 µL of resuspended beads and the appropriate quantity of respective target antibody (H3K9ac and H3K27ac) and IgG (control) was added, followed by incubation for 2 hours at 4°C, a brief spin on the magnetic separator for 1 minute. The supernatant was removed, and a cold SCW buffer containing Protease Inhibitor Cocktail III and 5 µL chromatin (final volume 500ul) was added (5 µL of undiluted chromatin was saved and used as 100% input) and rotated overnight at 4°C. The supernatant was removed and washed twice with 500 µL cold SCW buffer. The final wash was performed with 500 µL cold Low Stringency IP Wash Buffer containing Protease Inhibitor Cocktail III. Resuspend the beads in 500 µL cold Low Stringency Buffer containing Protease Inhibitor Cocktail III then transfer to a new microcentrifuge tube. Place the tube on the magnetic separator for 1 minute then remove the supernatant. For ChIP samples, resuspend beads in 50 µL ChIP Elution Buffer and add 1 µL Proteinase K, and to the 5 ul input chromatin, add 45 µL ChIP Elution Buffer and 1 µL Proteinase K. Incubate the samples in a Thermomixer at 65°C for 2 hours and then at 95°C for 15 minutes. Briefly spin the microcentrifuge tube briefly to settle contents; then place the tube on the magnetic separator for 1 minute. Carefully transfer 45 µL of the supernatant to a new microcentrifuge tube followed by q-PCR using the comparative Ct (ΔΔCt) method. q-PCR was performed to analyze acetylation of histone marks at the promoter of IFN-γ and TNFα using primer pairs of IFN-γ promoter (SimpleChIP Human IFN-γ Promoter Primers #13051) and TNFα promotor (forward sequence: GCC CCA GGG ACA TAT AAA GG; reverse sequence: GGC GTC TGA GGG TTG TTT T).

### Single-guide RNA (sgRNA)-mediated Gene Editing

Human Th1 cells were harvested, and CRISPR editing of human primary CD4^+^ T cells with ribonucleoprotein (RNP) complex using Nucleofector technology was done as per the manufacturer’s instructions. In brief, target-specific CRISPR SgRNAs and Cas9-NLS were purchased from Synthego. Lyophilized SgRNA was resuspended in Nuclease-free 1X TE buffer at a concentration of 250 pM and stored in aliquots at −20 °C. Nucleofector solution was prepared by adding a Supplement reagent to the P3 nucleofector solution (16.4 µl P3 Solution + 3.6 µl supplement per sample). RNP was prepared by incubating synthetic sgRNAs together with *Streptococcus pyogenes* Cas9 nuclease (SpCas9) at a ratio of 2.5:1. Incubate RNP mix at room temperature for 10 minutes. 1×10^6^ cells were washed twice with sterilized PBS and centrifuged at 1200 rpm for 5 minutes at room temperature. The supernatant was removed and resuspended in 20 µl of nucleofector solution. For each transfection, 20 µl of cell suspension was mixed with 5 µl of RNP solution and transferred into a well of the 16-well Nucleocuvette Strip. The cells were transfected using a Lonza 4D and X-unit electroporation system with pulse code EH-100. Immediately after electroporation, 100 µl of prewarmed medium was added to each well and the cells were incubated at 37 °C for 20 min. The cells were then transferred to a culture plate and cultured in complete RPMI 1640 with 10% FBS + anti-CD28 (2 ug/ml) for 5 days.

### Transcriptomic Analysis

Previously published single-cell RNA-seq data generated based on uninflamed and inflamed ileal resection specimens isolated from Crohn’s disease patients were analyzed as described earlier ^22^. Briefly, sequencing data were aligned to the human reference genome Grch38. Data with at least 500 unique molecular identifiers (UMIs), log10 genes per UMI >0.8, >250 genes per cell, and a mitochondrial ratio of less than 0.2% were extracted, normalized, and integrated using the Seurat package v3.0 in R 4.0.2. non-Treg CD4^+^ T cells (CD4^+^, FOXP3^-^, IL2RA^-^) were compared between inflamed and uninflamed samples as well as inflamed resection specimens of GIMATS (non-responders) and non-GIMATS (responders) datasets. Statistical differences were determined using an unpaired t-test.

### Statistical analysis

Please refer to the Figure legends for the description of sample size (*n*) and statistical details. Two-tailed unpaired Student’s t-test was used to compare the two groups. One-way ANOVA + Dunn’s or Bonferroni multiple comparisons test was used to compare three or more groups. Non-parametric murine data (DAI and MCHI) were analyzed using the Kruskal-Wallis test. *P* values less than 0.05 were considered statically significant. All data were analyzed using Prism (GraphPad Software, San Diego, CA).

**Supplementary Table 1:**
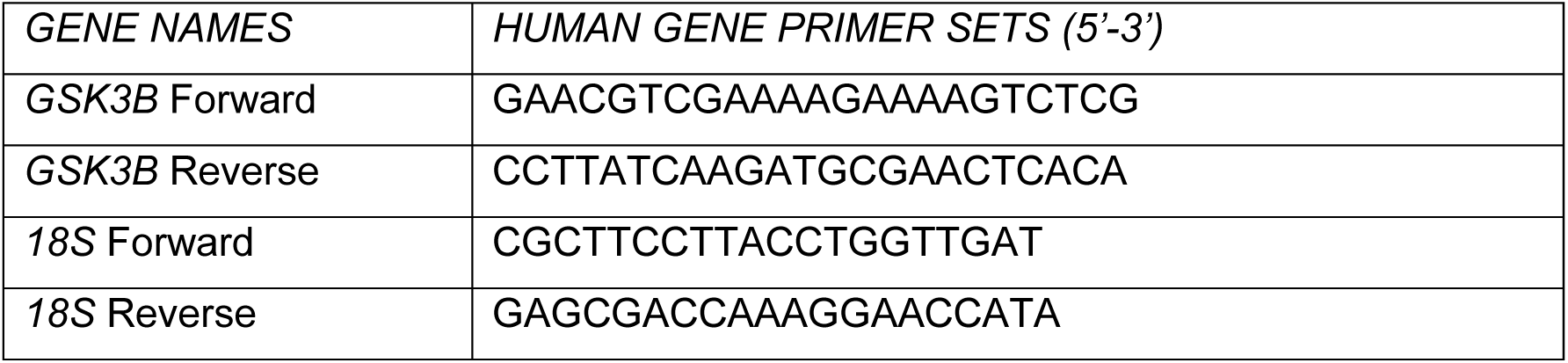
Primer sets for RT-qPCR

**Supplementary Table 2:**
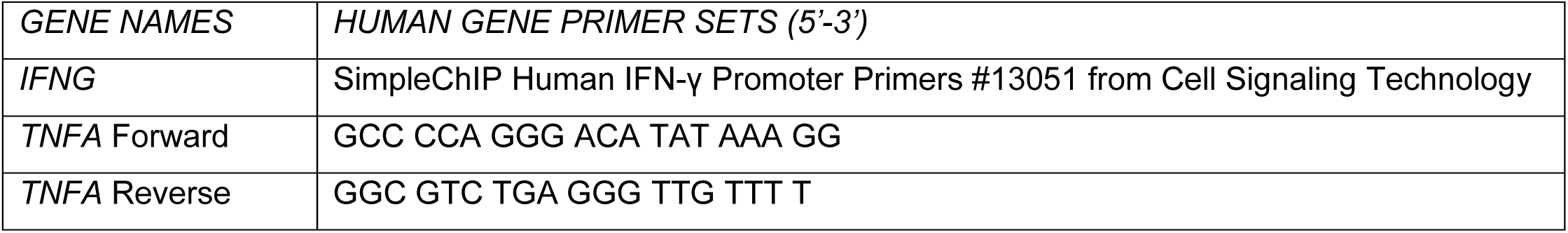
Primer sets for ChIP-qPCR

Key Resources Table

Key Resources Table
| REAGENT or RESOURCE | SOURCE | IDENTIFIER |
| --- | --- | --- |
| *Antibodies* |  |  |
| Anti-human/mouse/rat VDAC1 (Lot #3507477001) | ORIGENE | Cat#TA326858 |
| Anti-human/mouse/rat β-catenin | PetroTech | Cat#66379-1-Ig |
| Anti-human/mouse/rat/monkey phospho-Serine 216 GSK3β (D85E12) | Cell Signaling | Cat#5558S |
| Anti-phosphoserine | Millipore corp | Cat#05-1000 |
| Anti-human/mouse/rat/monkey GSK3β (27C10) | Cell Signaling | Cat#9315 |
| Anti-human/mouse/rat/monkey β-actin (8H10D10) | Cell Signaling | Cat#3700S |
| Anti-human/mouse/rat HK-I | Invitrogen | Cat#MA5-15680 |
| APC/Cyanine7 anti-human IFN-γ (4S.B3) | BioLegend | Cat#502530 |
| PE anti-human TNF-α (MAb11) | BioLegend | Cat#502909 |
| Alexa Fluor 488 anti-human CD4 (clone RPA-T4) | BioLegend | Cat#300519 |
| PE anti-mouse CD45RB (clone c363-16A) | BioLegend | Cat#103308 |
| Alexa Fluor 488 CD25 (clone PC61) | BioLegend | Cat#102017 |
| Alexa Fluor 488 Mouse IgG1, k (clone MOPC-21) | BioLegend | Cat#400134 |
| APC Mouse IgG1, k (clone MOPC-21) | BioLegend | Cat#400119 |
| PE Mouse IgG1, k (clone MOPC-21) | BioLegend | Cat#400114 |
| APC/Cyanine 7 Mouse IgG1, k (clone MOPC-21) | BioLegend | Cat#400128 |
| Brilliant Violet 421 Mouse IgG1, k (clone MOPC-21) | BioLegend | Cat#400157 |
| PE/Cy7 Mouse IgG1, k (clone MOPC-21) | BioLegend | Cat#400125 |
| PerCP/Cy5.5 Mouse IgG1, k (clone MOPC-21) | BioLegend | Cat#400150 |
| Human TruStain FcX™ | BioLegend | Cat#422302 |
| CellTak | Corning | Cat#354242 |
| *Chemicals, Peptides, and Recombinant Proteins* |  |  |
| LY2090314 | Selleckchem | Cat#S7063 |
| TCS 2002 | TOCRIS | Cat#3869 |
| Methyl Pyruvate | Sigma-Aldrich | Cat#371173 |
| MitoTracker Deep Red FM | Thermo Fisher Scientific | Cat#M46753 |
| Oligomycin | Abcam | Cat#ab141829 |
| FCCP | Abcam | Cat#ab120081 |
| Rotenone | Abcam | Cat#ab143135 |
| Antimycin A | Abcam | Cat#ab141904 |
| 2-deoxy-D-glucose (2-DG) | APEXBIO | Cat#B1027 |
| Seahorse XF 1.0 M Glucose solution | Agilent Technologies | Cat#103577-100 |
| Seahorse XF 200 mM Glutamine solution | Agilent Technologies | Cat#103579-100 |
| Seahorse XF 100 mM Pyruvate solution | Agilent Technologies | Cat#103578-100 |
| XF RPMI Medium | Agilent Technologies | Cat#103576-100 |
| Purified NA/LE Mouse Anti-human CD3 | BD Pharmingen | Cat#555329 |
| Purified NA/LE Mouse Anti-human CD28 | BD Pharmingen | Cat#555725 |
| Recombinant human IL-2 | PetroTech | Cat#200-02 |
| Recombinant human TGF-β1 | PetroTech | Cat#100-21C |
| Recombinant human IL-12 p70 | PetroTech | Cat#200-12 |
| Recombinant human IL-1β | PetroTech | Cat#200-01B |
| Recombinant human IL-6 | PetroTech | Cat#200-06 |
| Recombinant human IL-21 | PetroTech | Cat#200-21 |
| Recombinant human IL-23 | PetroTech | Cat#200-23 |
| Recombinant human IL-4 | PetroTech | Cat#200-04 |
| Purified anti-human IFN-γ (clone B27) | BioLegend | Cat#506502 |
| Purified anti-human IL-4 (clone 8D4-8) | BioLegend | Cat#500702 |
| Clotrimazole | TOCRIS | Cat#4096 |
| Fibronectin Human, Natural | Corning | Cat#354008 |
| Purified NA/LE Hamster Anti-mouse CD3e (clone145-2C11) | BD Pharmingen | Cat#553057 |
| Purified NA/LE Hamster Anti-mouse CD328 (clone145-2C11) | BD Pharmingen | Cat#553294 |
| Recombinant mouse IL-12 p70 | PetroTech | Cat#210-12 |
| Recombinant mouse IL-1β | PetroTech | Cat#211-11B |
| Recombinant mouse IL-6 | PetroTech | Cat#210-16 |
| Recombinant mouse IL-21 | PetroTech | Cat#210-21 |
| Recombinant mouse IL-23 p40 | R&D SYSTEMS | Cat#1887-ML-010 |
| LEAF<sup>TM</sup> Purified anti-mouse IFN-γ (clone XMG1.2) | BioLegend | Cat#505812 |
| Purified anti-mouse IL-4 (clone 11B11) | BioLegend | Cat#504102 |
| *Biological Samples and Experimental Models* |  |  |
| Healthy adult – peripheral blood mononuclear cells (males) | Mayo Clinic | N/A |
| C57BL/6J mice (males) | The Jackson Laboratory | Stock no:000664 |
| C57BL/6J *Rag1*<sup>-/-</sup> mice (males) | The Jackson Laboratory | Stock no: 002216 |
| C57BL/6NJ wildtype (one male and one female) | The Jackson Laboratory | Stock no: 005304 |
| Total body *Il21r*<sup>-/-</sup> (one male and one female) | The Jackson Laboratory | Stock no: 019115 |
| RAG1<sup>-/-</sup> mice (male) | The Jackson Laboratory | Stock no: 002216 |
| *Software and Algorithms* |  |  |
| FlowJo version10.2 | FlowJo, LLC | https://www.flowjo.com |
| Zen 3.0 (blue edition) | Zen | https://www.zeiss.com |
| Bio-Rad Image Lab | Bio-Rad | https://www.bio-rad.com |
| GraphPadPrism 8 | GraphPad software | https://www.graphpad.com |
| Seahorse Bioscience Wave Desktop | Agilent Technologies | https://www.agilent.com |
| ImageJ | NIH | https://imagej.nih.gov |
| Photoshop Adobe | Adobe | https://www.adobe.com |
| *Critical Commercial Assays* |  |  |
| BD<sup>TM</sup> Cytometric Bead Array (CBA) Mouse Th1/Th2/Th17 Cytokine Kit | BD Biosciences | Cat#560485 |
| RNeasy Plus Mini Kit | QIAGEN | Cat#74134 |
| Seahorse XFe96 Flux/Pak | Agilent Technologies | Cat#102416-100 |
| Duolink™ In Situ PLA Anti-Mouse PLUS | Sigma-Aldrich | Cat#DUO92001 |
| Duolink™ In Situ PLA Anti-Rabbit MINUS | Sigma-Aldrich | Cat#DUO92005 |
| Duolink™ In Situ Detection Reagents Red | Sigma-Aldrich | Cat#DUO92008 |
| CD4<sup>+</sup> T cell isolation kit, mouse | Miltenyi Biotec | Cat#130-104-454 |
| CD4<sup>+</sup> CD62L<sup>+</sup> (naïve) T cell isolation kit, mouse | Miltenyi Biotec | Cat#130-106-643 |
| CD4<sup>+</sup> T cell isolation kit, human | Miltenyi Biotec | Cat#130-096-533 |
| CD45RA<sup>+</sup> (naïve) cell isolation kit, human | Miltenyi Biotec | Cat#130-045-901 |
| NanoString Metabolism Panel | NanoString technology | Cat#115000367 |
| NanoString Autoimmune Panel | NanoString technology | Cat#115000262 |
| Negative control scrambled sgRNA#1 | Synthego Corporation | CRISPR Revolution sgRNA EZ Kit |
| Human *GSK3B* sgRNA | Synthego Corporation | Cat#120002213 |
| SpCas9, 1000pmol | Synthego Corporation | SpCas9, 1000pmol |
| P3 Primary Cell 4D X Kit | Lonza Biosciences | Cat#V4XP-3032 |
| *Others* |  |  |
| RPMI Medium 1640 (1X) [+] L-Glutamine | Gibco | REF#11875-093 |

## Results

### Human Effector Th1 Cells Display High Aerobic Glycolysis but Low Mitochondrial Respiration

To investigate the metabolic underpinnings of CD4^+^ T cell inflammatory response, we performed real-time metabolic phenotyping of human Th1 cells via Seahorse analysis. Naïve CD4^+^ T cells isolated from human peripheral blood mononuclear cells (PBMCs) of healthy donors were differentiated *in vitro* into Th1, Th2, and Th17 cells *vs*. functionally distinct induced Tregs (iTregs) (Figure 1A). Cells were subjected to mitochondrial and glycolysis stress tests to assess mitochondrial respiration (i.e., oxygen consumption rate, OCR) and glycolytic function (i.e., extracellular acidification rate, ECAR). Seahorse analysis revealed that basal, ATP-linked and maximal respiration and spare respiratory capacity were low in effector T cells, particularly in Th1 cells, compared to iTregs (Figures 1B and 1C). In contrast, effector Th1 and Th17 cells exhibited high ECAR from glucose-induced glycolysis and oligomycin-induced glycolytic capacity compared to iTregs (Figures 1D and 1E). Therefore, consistent with previous reports^23^, in response to glucose catabolism, effector Th1 cells are highly glycolytic at steady state but less capable of mitochondrial respiration.

**Figure 1.**
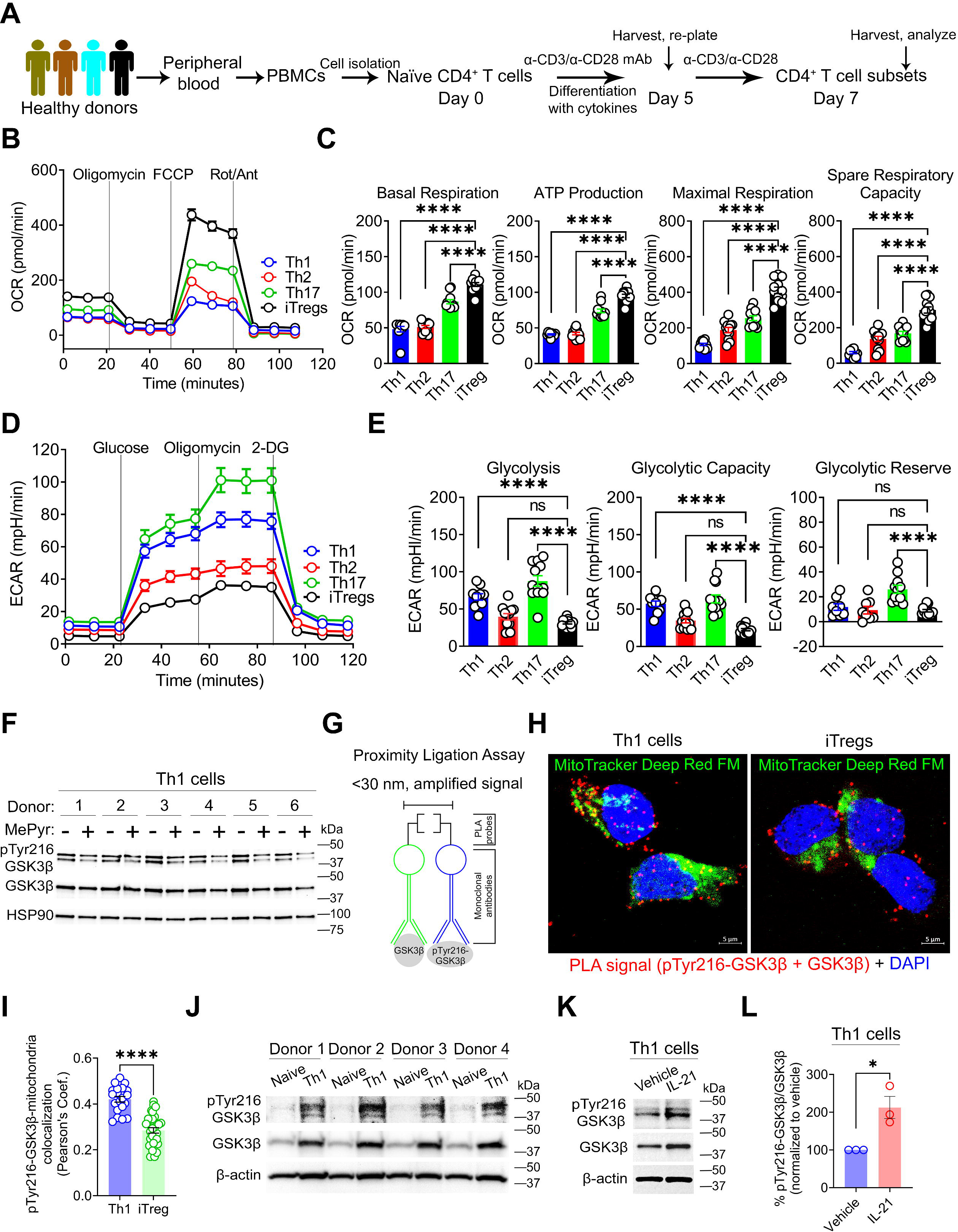
Human Effector Th1 Cells Display High Aerobic Glycolysis but Low Mitochondrial Respiration. (A) Experimental workflow for naïve CD4^+^ T cell isolation from PBMCs and polarization into CD4^+^ T cell subsets. (B and C) Representative OCR profile of human CD4^+^ T cell subsets before and after mitochondrial perturbation (n = 3 biological replicates) (B). The bar graphs show calculated basal respiration, ATP production (ATP-linked), maximal respiration, and spare respiratory capacity; mean ± SEM from 10-12 technical replicates (C). (D and E) Representative ECAR profile of human CD4^+^ T cell subsets (n = 3 biological replicates) (D). The bar graphs show calculated glycolysis, glycolytic capacity, and glycolytic reserve; mean ± SEM from 10-12 technical replicates (E) (F) Immunoblot analysis of whole cell lysates (WCLs) derived from human effector Th1 cells shows active GSK3β protein expression relative to total GSK3β and HSP90 ± MePyr 5 mM for 24 h (n = 6 biological replicates). (G) Schematic of proximity ligation assay (PLA), which generates a positive PLA signal for the detection of active GSK3β (pTyr216-GSK3β) if the distance between GSK3β and pTyr216-GSK3β primary monoclonal antibodies bound to the same GSK3β protein is less than 30 nm (<30 nm). (H and I) Representative confocal images of human effector Th1 cells *vs*. iTregs show pTyr216-GSK3β protein expression and localization in red after PLA, mitochondrial staining using MitoTracker Deep Red FM in pseudo color green, and nuclei staining using 4’,6’-diamidino-2-phenylinode (DAPI) (DNA, blue); scale bar, 5 µm (H). The graph shows Pearson’s Coefficient quantifying the colocalization of PLA signals (pTyr216-GSK3β) with mitochondria (n = 20 Th1 cells, n = 35 iTregs) (I). (J) Immunoblot analysis of WCLs shows pTyr216-GSK3β protein expression relative to the total GSK3β and β-actin in human naïve cells *vs*. effector Th1 cells. (K and L) Representative immunoblot analysis of WCLs shows pTyr216-GSK3β protein expression relative to the total GSK3β and β-actin in human effector Th1 cells ± IL-21 100 ng/ml for 24 h (K). The bar graph shows the quantitation of pTyr216-GSK3β protein expression relative to the total GSK3β. Data represents mean ± SEM. * p < 0.05, ** p < 0.01, *** p < 0.001, and **** p < 0.0001, using two-tailed Student’s t-test or one-way ANOVA followed by Bonferroni test for multiple comparisons.

Glucose-derived pyruvate lies at the bifurcation point since it’s either converted to lactate or metabolized in the TCA cycle to drive OXPHOS. To reveal the intrinsic intracellular events resisting OXPHOS, pyruvate metabolism was enforced by supplementing effector Th1 cells with a mitochondria membrane-permeable form of pyruvate (methyl pyruvate, MePyr) for 24 h, followed by assessment of GSK3β, a serine-threonine kinase reported to inhibit mitochondrial pyruvate metabolism and drive glycolysis in non-CD4^+^ T cells^24–27^. MePyr had no effect on *GSK3B* mRNA (Supplementary Figure 1A); however, activated GSK3β (phosphorylated GSK3β on Tyrosine residue 216, pTyr216-GSK3β) was reduced (Figure 1F). This result suggests that MePyr may exert its metabolic effect on effector Th1 cells by suppressing GSK3β enzymatic activity.

We then speculated that pTyr216-GSK3β may localize to the mitochondria to control pyruvate entry. MitoTracker Deep Red FM dye staining of live effector Th1 cells (shown as the pseudo color green) and subsequent Proximity Ligation Assay (PLA) using monoclonal antibodies against pTyr216-GSK3β and pTyr216-GSK3β (Figure 1G) revealed that mitochondrial pTyr216-GSK3β (which refers to pTyr216-GSK3β in proximity to the mitochondria as evidenced by the discrete red PLA signals in colocalization with the pseudo color green; Pearson’s Coefficient) was higher in effector Th1 cells than iTregs (Figures 1H and 1I). Investigating the extracellular signals governing GSK3β expression, immunoblot analyses revealed that GSK3β and pTyr216-GSK3β protein expression were elevated in effector Th1 cells compared to control naïve CD4^+^ T cells (Figure 1J), implying that T cell receptor (TCR) and CD28 ligation promotes GSK3β expression and phosphorylation. Furthermore, stimulation with IL-21, a proinflammatory cytokine associated with immune-mediated diseases^28^ and recently shown to activate GSK3β in Tregs^29^, increased pTyr216-GSK3β levels (Figures 1K and 1L). Thus, guided by microenvironmental cues, GSK3β may serve as a metabolic checkpoint mediating glucose conversion to lactate rather than its utilization for OXPHOS, thereby sustaining the Th1 cell inflammatory program.

### MePyr Supplementation Induces Effector Th1 Cell Transcriptional Rewiring via GSK3β Inhibition

Next, we explored whether MePyr can alter the cellular state of Th1 cells in a manner mirroring direct pharmacologic inhibition of GSK3β. To obtain a comprehensive dataset defining the Th1 cell transcriptional state, RNA isolated from human effector Th1 cells supplemented with or without MePyr for 24 h were subjected to NanoString-based mRNA profiling assays that quantified the copy numbers of 786 metabolism and 770 autoimmune-related human genes (Figure 2A). For mRNA profiling using the metabolism panel, 514 genes were detected (Figure 2B), 41 of which were upregulated in MePyr-supplemented *vs*. untreated Th1 cells (including Treg and OXPHOS-related genes; *FOXP3*, *ICOS*, *TIGIT*, *NDUFB8*, *IDH3A*, *IDH3B*, and *NQO1*) (Supplementary Figure 2A, top left), while 6 genes were downregulated (Supplementary Figure 2A, bottom left). To assess similarities between transcriptional changes induced by MePyr supplementation and GSK3β inhibition with LY2090314, an inhibitor deemed safe in a phase 2 study in patients with acute leukemia^30^, we performed NanoString-based mRNA profiling of LY2090314 *vs*. DMSO for 24 h (Figure 2C). 514 genes were detected (Figure 2C), 84 of which were upregulated in LY2090314-treated Th1 cells *vs*. DMSO (including Treg and OXPHOS-related genes; *GZMA*, *GZMB*, *SMAD2*, *ICOS*, *TIGIT, ACAA2, NDUFA12, NDUFA1,* and *IDH1*) (Supplementary Figure 2A, top right), while 24 genes were downregulated (such as Th cell and glycolysis-related genes; *TBX21, MYC, IL21R, RRM2, STAT1, ENO1,* and *TNF*) (Supplementary Figure 2A, bottom right). The 16 significant, overlapping upregulated genes in both MePyr-supplemented *vs*. LY2090314-treated Th1 cells include Treg-related genes, such as *IL10*, *GZMH*, *ASS1*, and *CTLA4*) (Supplementary Figure 2A, top), while only *COL6A3* and *EHHADH* were downregulated (Supplementary Figure 2A, bottom). Gene Ontology (GO) biological process pathway analysis of the 16 overlapping upregulated genes in MePyr and LY2090314-treated cells revealed an association with pathways involved in the negative regulation of immune effector processes, leukocyte activation, and immune response (Supplementary Figure 2B). Furthermore, KEGG pathway analysis of these 16 upregulated overlapping genes revealed an association with processes involved in the regulation of immune-mediated diseases (such as IBD), metabolic pathways, TCR, IL-17, JAK-STAT and HIF-1 signaling (Supplementary Figure 2C). Therefore, in a manner mirroring GSK3β inhibition, MePyr supplementation transcriptionally rewires effector Th1 cells toward a Treg-like metabolic program (Figures 1B-1D) as evidenced by the enhanced expression of OXPHOS-related genes such as the mitochondrial NADH dehydrogenase subcomplexes (*NDUFB8, NDUFA12,* and *NDUFA1*) encoding subunits of the largest respiratory complex, the electron transport complex 1 (ETC1).

**Figure 2.**
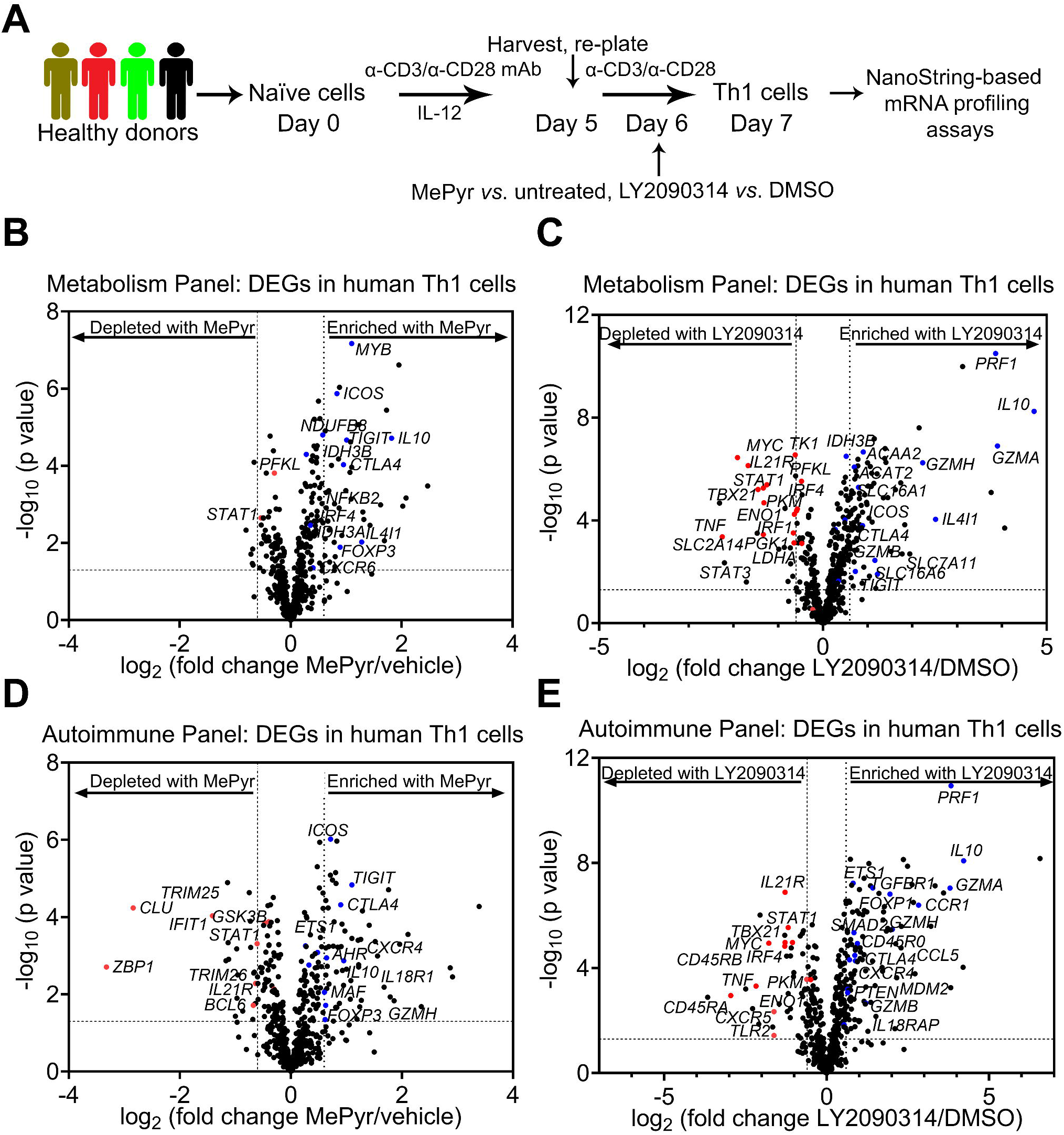
Methyl Pyruvate Supplementation Induces Effector Th1 Cell Transcriptional Rewiring via GSK3β Inhibition. (A) Experimental workflow for human effector Th1 cell supplementation with methyl pyruvate (MePyr 5 mM) *vs*. vehicle (DMSO) or treatment with GSK3β inhibitor (LY2090314 0.0125 µM) *vs*. DMSO for 24 h, followed by NanoString-based mRNA profiling using the NanoString nCounter System. (B and C) The volcano plots illustrate the expression of all 514 detected genes in effector Th1 cells (MePyr 5 mM /untreated 24 h, n = 4 biological replicates) (B) or effector Th1 cells (LY2090314 0.0125 µM/DMSO, n = 4 biological replicates) (C) using the NanoString Metabolism Panel. Red and blue dots denote relevant downregulated and upregulated genes within the differentially expressed genes (DEGs), respectively. (D and E) The volcano plots illustrate the expression of all 558 detected genes in effector Th1 cells (MePyr 5 mM/untreated, n = 4 biological replicates) or effector Th1 cells (LY2090314 0.0125 µM/DMSO, n = 4 biological replicates) using the NanoString Autoimmune Panel 24 h after treatment. Red and blue dots denote relevant downregulated and upregulated genes within the DEGs, respectively.

For mRNA profiling of MePyr-supplemented *vs*. untreated Th1 cells using the autoimmune panel, 558 genes were detected (Figure 2D), 58 (including Treg-related genes; *FOXP3*, *IL10, GZMH, ICOS, AHR, TNFRSF9, CTLA4, TIGIT*) and 23 (including *STAT1* and *ITGB7*) of which were upregulated and downregulated, respectively (Supplementary Figure 2D, top left and bottom left). Similarly, LY2090314 treatment upregulated 110 genes (including Treg-related genes; *TGBR1, FOXP1, GZMA, GZMB, SMAD2, MAF, CTLA4, TIGIT* as shown in Figures 2E and Supplementary 2D, top right) and downregulated 43 genes (such as glycolysis and Th cell-related genes; *IL21R, TBX21, ENO1, TNF*) *vs*. DMSO-treated cells (Supplementary Figure 2D, bottom right). The 25 genes significantly upregulated in both MePyr-supplemented cells vs. LY2090314-treated cells include Treg-related genes, such as *IL10, GZMH, TIGIT, TNFRSF9, CTLA4* (Supplementary Figure 2D, top), while *STAT1, ITGB7, RARRES3,* and *CLU* genes were downregulated (Supplementary Figure 2D, bottom). GO biological process pathway analysis of the 25 overlapping upregulated genes revealed an association with pathways involved in the regulation of cytokine production and cell activation, proliferation, and adhesion (Supplementary Figure 2E). Moreover, KEGG pathway analysis of these same upregulated overlapping genes revealed an association with processes involved in regulating immune-mediated diseases (such as IBD) and IL-17, TCR and JAK-STAT signaling (Supplementary Figure 2F). Therefore, mirroring GSK3β inhibition, MePyr transcriptionally rewires inflammatory Th1 cells toward a Treg-like phenotypic program. Overall, via NanoString-based mRNA profiling of metabolic and autoimmune-related genes, MePyr or GSK3β inhibition transcriptionally induces glycolysis to OXPHOS metabolism in Th1 cells with concomitant loss of inflammatory phenotype and gain of Treg-like program.

### MePyr Supplementation Induces Metabolic Rewiring of Effector Th1 Cells by Suppressing Glycolysis

To define how MePyr induced transcriptional alterations, we subjected MePyr-treated effector Th1 cells to a glycolysis stress test (Figure 3A). Effector Th17 cells were similarly profiled, given that these inflammatory cells are also highly glycolytic (Figure 1D). In support of NanoString-based mRNA profiling assays, MePyr reduced ECAR associated with basal glycolysis and glycolytic capacity of effector Th1 cells (Figures 3B and 3C) and Th17 cells (Supplementary Figures 3A-3C). Similarly, LY2090314 treatment mirrored MePyr supplementation as evidenced by reduced ECAR associated with basal glycolysis, glycolytic capacity, and glycolytic reserve in Th1 cells (Figures 3D and 3E) and Th17 cells (Supplementary Figures 3D and 3E). These suggest that MePyr supplementation of effector Th1 and Th17 cells impairs glucose catabolism to lactate via GSK3β inhibition.

**Figure 3.**
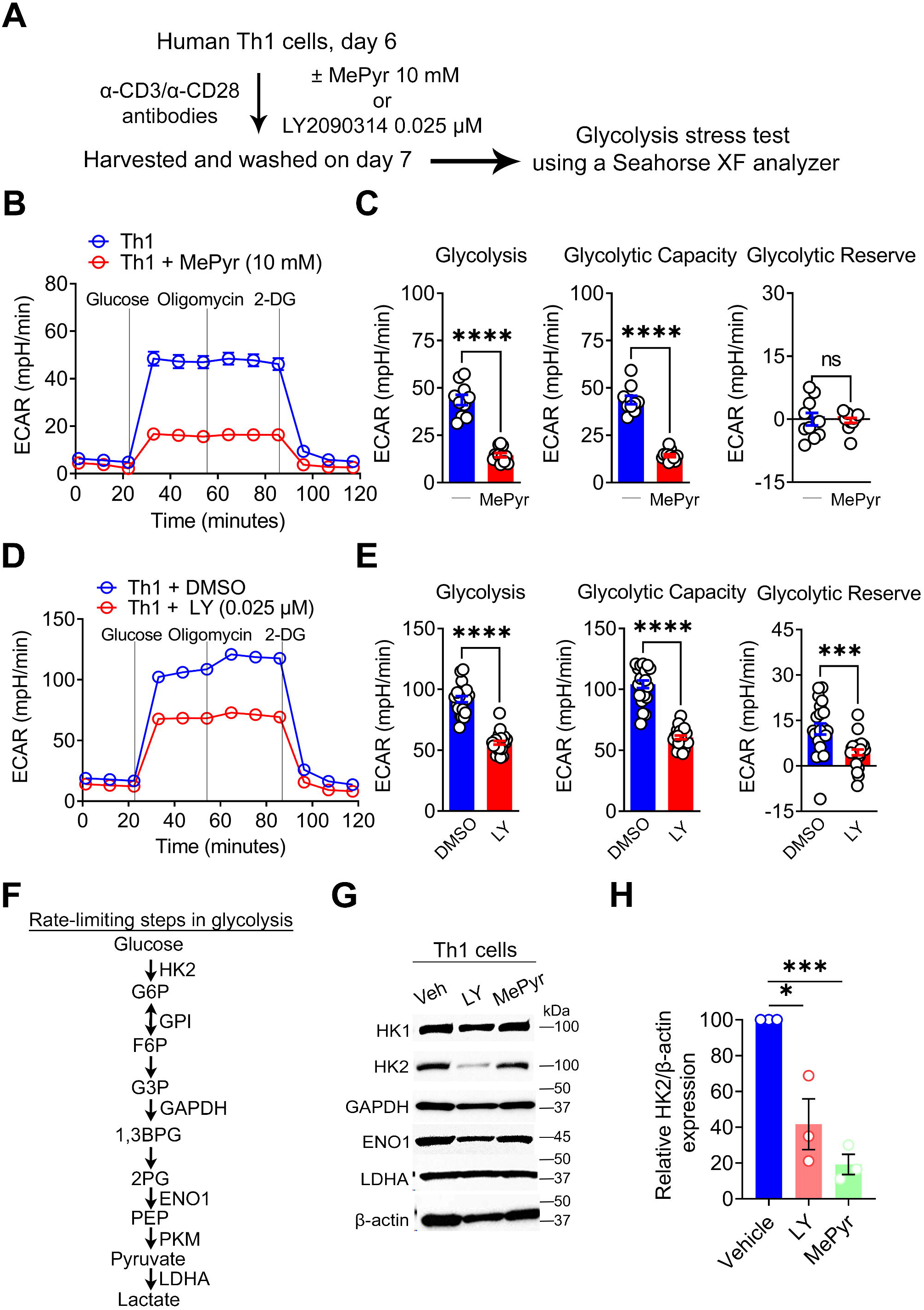
Methyl Pyruvate Induces Metabolic Rewiring of Human Effector Th1 Cells. (A) Experimental workflow for human effector Th1 cell supplementation with MePyr 10 mM *vs*. untreated or treatment with LY2090314 0.025 µM *vs*. DMSO for 36 h, followed by Seahorse analysis. (B and C) Representative ECAR profile of human effector Th1 cells 36 h after supplementation with MePyr 10 mM compared to untreated cells (n = 3 biological replicates) (B). The bar graphs show calculated glycolysis, glycolytic capacity, and glycolytic reserve; mean ± SEM from 10-12 technical replicates (C). (D and E) Representative ECAR profile of human effector Th1 cells 36 h after supplementation with LY2090314 (LY) 0.025 µM compared to DMSO-treated cells (n = 3 biological replicates) (D). The bar graphs show calculated glycolysis (basal), glycolytic capacity, and glycolytic reserve; mean ± SEM from 10-12 technical replicates (E). (F) Schematic illustrates glycolytic steps and enzymes involved in glucose catabolism to lactate. (G and H) Representative immunoblots show the protein expression of glycolytic enzymes in WCLs derived from human effector Th1 cells after treatment with MePyr 10 mM or LY2090314 (LY) 0.025 µM for 36 h (G). The graph shows quantitation of hexokinase 2 (HK2) protein expression relative to β-actin (H). Data represents mean ± SEM. * p < 0.05, *** p < 0.001, and **** p < 0.0001 using a two-tailed Student’s t-test. ns (not significant).

To illuminate how MePyr impaired glycolysis, we measured the protein expression of key rate-limiting enzymes involved in glucose catabolism and aerobic glycolysis (Figure 3F), such as HK2; hexokinase 2, GAPDH; glyceraldehyde-3-phosphate dehydrogenase, ENO1; enolase 1, and LDHA; lactate dehydrogenase. HK2 is the first and rate-limiting enzyme in the glycolytic pathway that catalyzes glucose conversion to glucose-6-phosphate (G6P), ENO1 catalyzes 2-phosphoglycerate (2-PG) conversion to phosphoenolpyruvate (PEP), and LDHA and LDHB, which forms five tetrameric LDH isoenzymes and defines the biochemical reaction of aerobic glycolysis by converting pyruvate to lactate. Like LY2090314, MePyr reduced HK2 expression, while GAPDH, ENO1, and LDHA levels were unaffected (Figures 3G and 3H). Overall, MePyr-mediated enforcement of mitochondrial metabolism attenuates HK2-induced glycolysis by deactivating GSK3β kinase activity.

### Methyl Pyruvate Suppresses Inflammatory Cytokine Production by Effector Th1 Cells via Attenuation of Histone Acetylation

Aerobic glycolysis is a well-documented driver of Th1 and Th17 cell immune responses, including cytokine production^14, 15, 20, 23^. Given that MePyr attenuated HK2-mediated glycolysis, we postulated this metabolic event would weaken inflammatory cytokine production by Th1 cells. As expected, MePyr reduced the percentage of IFN-γ^+^ and TNF-α^+^ Th1 cells *vs*. vehicle-treated cells (Figures 4A-4C). Moreover, consistent with NanoString-based mRNA profiling in which *TNF* was reduced by LY2090314 (Figures 2C and 2E), GSK3β inhibitors (LY2090314 and TCS2002) reduced the percentage of IFN-γ^+^ and TNF-α^+^ Th1 cells *vs*. vehicle-treated cells (Figures 4A-4C). As expected, Th1 cells treated with 2-deoxy-Ɒ-glucose (2-DG), a well-documented inhibitor of HK2-mediated glycolysis, reduced the percentage of IFN-γ^+^ and TNF-α^+^ Th1 cells (Supplementary Figures 4A and 4B). This indicates that enforced mitochondrial pyruvate metabolism in effector Th1 cells limits GSK3β-HK2-mediated glycolysis, resulting in impaired production of IFN-γ and TNF-α.

**Figure 4.**
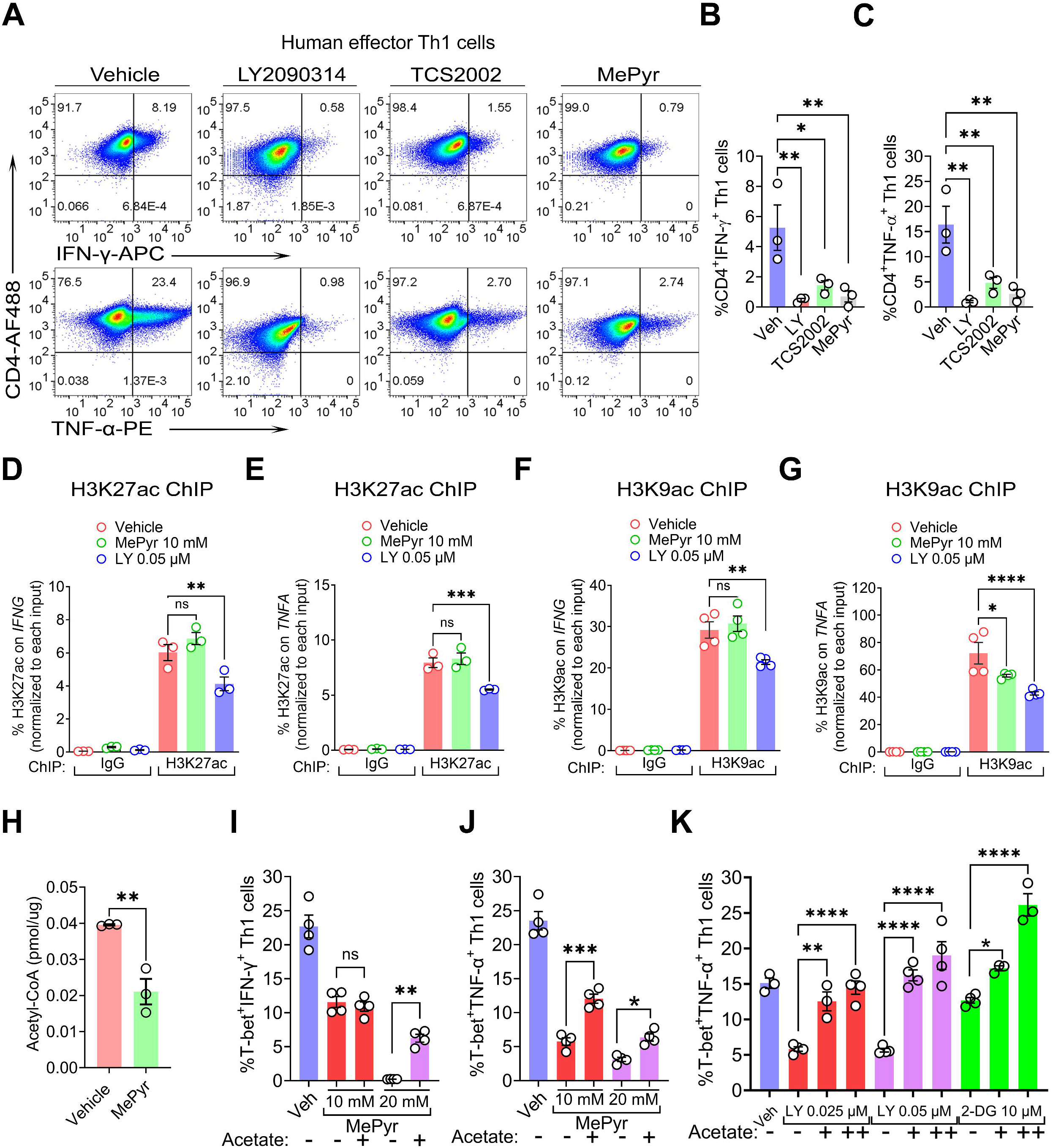
Methyl Pyruvate Suppresses Inflammatory Cytokine Production by Effector Th1 Cells via Attenuation of Histone Acetylation. (A-C) Dot plots indicate the percentage of human CD4^+^IFN-γ^+^ (double-positive) effector Th1 cells (top panels) or CD4^+^TNF-α^+^ effector Th1 cells (bottom panels) as determined via FACS analysis after exposure to LY2090314 (LY 0.05µM), TCS2002 0.5 µM or MePyr 10 mM for 36 h. The bar graphs show the percentage of CD4^+^IFN-γ^+^ effector Th1 cells (B) or CD4^+^TNF-α^+^ effector Th1 cells (C) (n = 3 biological replicates). (D and E) The bar graphs show the H3K27ac enrichment on the *IFNG* (D) and *TNFA* (E) promoters in human effector Th1 cells ± MePyr or LY2090314 (LY) for 36 h as determined via H3K27ac ChIP-qPCR (n = 3-4 biological replicates). (F and G) The bar graphs indicate H3K9ac enrichment on the *IFNG* (F) and *TNFA* (G) promoters in human effector Th1 cells ± MePyr or LY2090314 (LY) for 36 as determined via H3K27ac ChIP-qPCR (n = 3-4 biological replicates). (H) The gar graphs show intracellular acetyl-CoA levels in human effector Th1 cells after supplementation with MePyr 10 mM for 36 h. (I and J) The bar graphs show the percentage of human T-bet^+^IFN-γ^+^ effector Th1 cells (I) or T-bet^+^TNF-α^+^ effector Th1 cells (J) ± MePyr, sodium acetate 50 mM for 36 h as determined via FACS analysis (n = 3-4 biological replicates). (K) The bar graphs show the percentage of human T-bet^+^TNF-α^+^ effector Th1 cells ± LY2090314 (LY), 2-DG, sodium acetate 20-50 mM for 36 h as determined via FACS analysis (n = 3-4 biological replicates). Data represents mean ± SEM. * p < 0.05, ** p < 0.01, *** p < 0.001 and **** p < 0.0001, using two-tailed Student’s t-test or one-way ANOVA followed by Bonferroni test for multiple comparisons.

Glucose metabolism controls gene expression via epigenetic processes, such as histone H3 acetylation of lysine 9 and 27 residues^31, 32^. Although *IFNG* mRNA expression was not reduced by MePyr supplementation or LY2090314 treatment (Figures 2F and 2G), NanoString-based mRNA profiling revealed a significant reduction of *TNFA* mRNA by LY2090314 (Figures 2C and 2E). The discrepancy between *IFNG* and *TNFA* mRNA regulation may be attributed to experimental or biological technicalities. Therefore, we assessed whether the reduction in cytokine mRNA and the frequency of cytokine-producing Th1 cells in response to MePyr or GSK3β inhibition was due to the weakened presence of acetylated histones on *TNFA* and *IFNG* promoter regions. We performed chromatin immunoprecipitation-qPCR (ChIP-qPCR) analysis of acetylated histone H3 at lysine 27 and 9 residues (H3K27ac and H3K9ac) given that these histone marks are associated with active gene transcription. H3K27ac-ChIP and subsequent qPCR analyses revealed a decrease in the H3K27ac mark on *IFNG* and *TNFA* promoters in LY2090314-treated Th1 cells but not in MePyr-supplemented cells (Figures 4D and 4E, respectively). LY2090314 reduced the H3K9ac mark on the *IFNG* promoter, which was not observed in MePyr-supplemented cells (Figure 4F); however, both MePyr and LY2090314 diminished the H3K9ac mark on the *TNFA* promoter (Figure 4G). Therefore, MePyr supplementation or GSK3β inhibition abrogates IFN-γ and TNF-α expression by potentially diminishing acetylated histone H3 enrichment on cytokine loci. This is consistent with a study in which LDHA-mediated glycolysis maintained high cytoplasmic acetyl-coenzyme A (acetyl-CoA) concentration needed for histone acetylation and IFN-γ transcription in Th1 cells via glucose-driven mitochondrial acetyl-CoA sustaining citrate to acetyl-CoA shuttle system in the cytoplasm^19^.

Histone acetylation requires acetyl-CoA as a substrate, and glucose is a critical source of mitochondrial acetyl-CoA, which consequently sustains citrate to acetyl-CoA shuttle system required for histone H3 acetylation and IFN-γ transcription^19^. Considering the heightened OXPHOS (Figures 2B-2G) and glycolytic deficit (Figures 3B-3G) displayed by MePyr-supplemented and LY2090314-treated Th1 cells, we hypothesized that acetyl-CoA deficiency in these cells likely contributes to reduced histone H3 acetylation and cytokine expression. Indeed, we found that MePyr supplementation of Th1 cells reduced the accumulation of acetyl-CoA (Figure 4H). To demonstrate that impaired cytokine production was due to acetyl-CoA deficiency, we assessed whether exogenous supplementation of acetate could restore cytokine levels. Acetyl-CoA can be derived from acetate by acetyl-CoA synthetase (ACSS)-dependent ligation of acetate and CoA independent of citrate^33^. Sodium acetate (Na acetate) supplementation partially restored the frequency of IFN-γ^+^ and TNF-α^+^ populations as well as IFN-γ and TNF-α expression in MePyr-supplemented Th1 cells (Figures 4I, 4J, and Supplementary Figure 4C) and LY2090314-treated Th1 cells (Figure 4K and Supplementary Figures 4D, 4E). We further reasoned that 2-DG-treated Th1 cells supplemented with acetate would circumvent the need for glycolysis linked to immune response. As expected, the acetate supplementation of 2-DG-treated Th1 cells restored IFN-γ^+^ and TNF-α^+^ populations and expression of IFN-γ and TNF-α (Figure 4K and Supplementary Figures 4D, 4E). Overall, MePyr or GSK3β inhibition impairs acetylated histone H3 enrichment, resulting in the attenuation of IFN-γ and TNF-α levels in a manner reversible by acetate supplementation.

### IL-21-induced GSK3β activation and Metabolic Rewiring Sustains Inflammatory Response by Th1 Cells

CD4^+^T cells have been previously described to express IL-21^20, 34–36^. Given the observation that GSK3β is active in effector Th1 cells (Figure 1J), which was further enhanced by IL-21 (Figures 1K and 1L), we speculated that the autocrine IL-21-IL-21R signaling sustains GSK3β activity to augment glycolysis in Th1 and Th17 cells. Effector Th1 and Th17 cells, and iTregs (via naïve CD4^+^ T cell polarization) from WT *vs*. IL-21R knockout (*Il21r*^-/-^) mice^37^ were subjected to a glycolysis stress test. Notably, *Il21r*^-/-^ Th1 cells (Figures 5A and 5B) exhibited reduced basal glycolysis, glycolytic capacity, and glycolytic reserve compared to WT Th1 cells, while *Il21r*^-/-^ Th17 cells exhibited reduced glycolytic capacity compared to WT Th17 cells (Supplementary Figures 5A and 5B). In contrast to Th1 and Th17 cells, there was no difference in ECAR between WT iTregs *vs*. *Il21r*^-/-^ iTregs (Supplementary Figures 5C and 5D). Therefore, in the absence of exogenous IL-21, IL-21R deficiency in effector Th1 and Th17 cells is sufficient to impair glycolysis.

**Figure 5.**
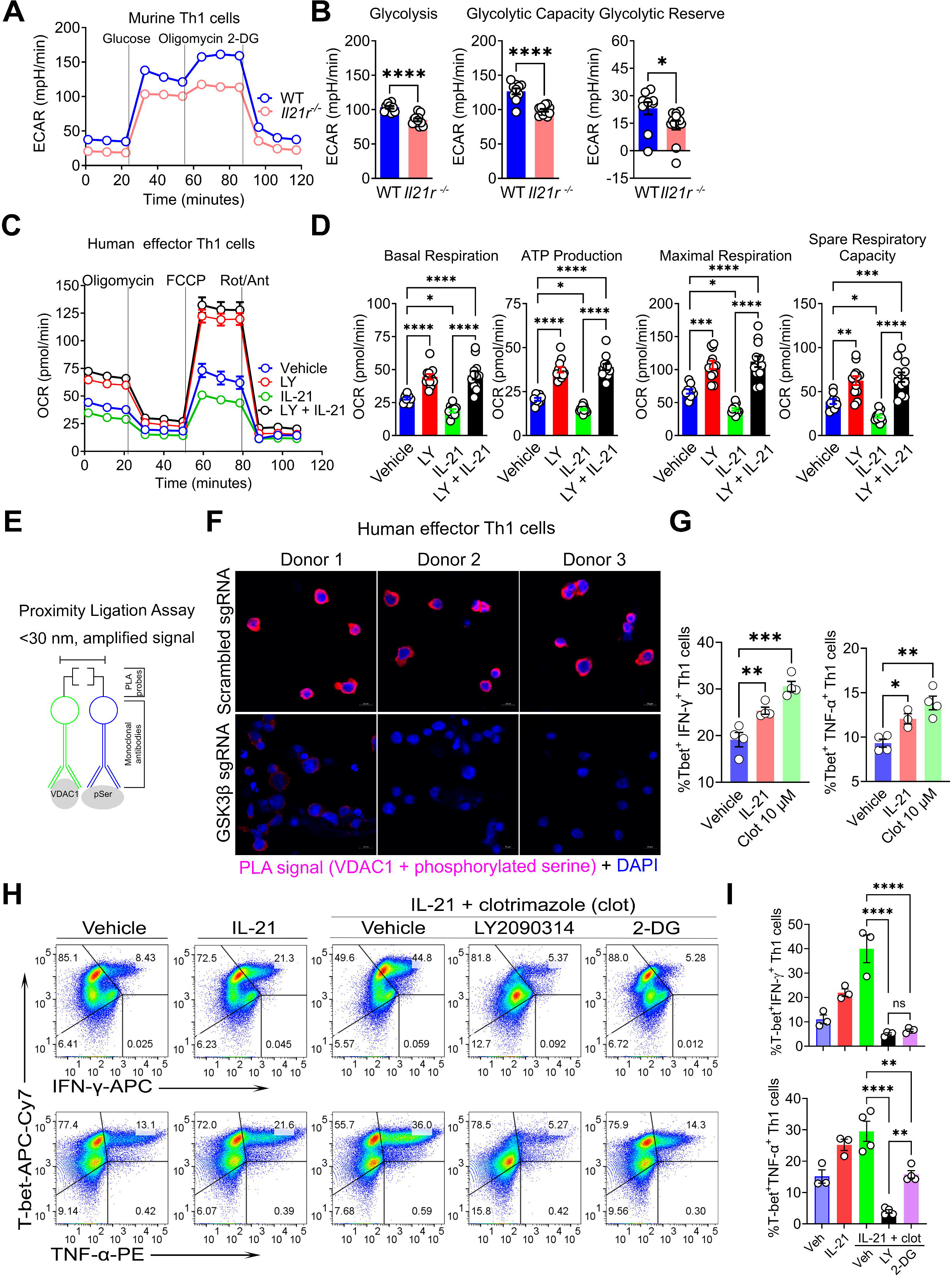
IL-21-induced GSK3β activation and Metabolic Rewiring sustain inflammatory Response by Th1 Cells. (A and B) Representative ECAR profile of differentiated WT *vs*. IL-21R^-/-^ effector Th1 cells from mice (cells were pooled from n = 5 mice per group) (A). The bar graphs show calculated glycolysis, glycolytic capacity, and glycolytic reserve; mean ± SEM from 10-12 technical replicates (B). (C and D) Representative OCR profile of human effector Th1 cells (treated with vehicle, LY2090314 (LY), IL-21 or IL-21 and LY for 36 h) before and after mitochondrial perturbation (n = 3 biological replicates) (C). The bar graphs show calculated basal respiration, ATP production, maximal respiration, and spare respiratory capacity; mean ± SEM from 10-12 technical replicates (D). (E) Schematic of PLA experiment, which generates a positive PLA signal for the detection of phosphorylated serine on VDAC1 (pSer-VDAC1) if the distance between phosphorylated serine and VDAC1 primary monoclonal antibodies bound to the same VDAC1 protein is less than 30 nm (<30 nm). (F) Representative confocal images of human effector Th1 cells nucleofected with scrambled sgRNA or GSK3β sgRNA show pSer-VDAC1 protein expression in red after PLA and nuclei staining using DAPI (DNA, blue); scale bar, 10 µm. (G) The bar graphs show the percentage of human T-bet^+^IFN-γ^+^ effector Th1 cells (left) or T-bet^+^TNF-α^+^ effector Th1 cells (right) after exposure to the vehicle (DMSO), IL-21+DMSO, or clotrimazole (clot) for 36 h as determined via FACS analysis (n = 4 biological replicates). (H and I) Dot plots show the percentage of T-bet^+^IFN-γ^+^ effector Th1 cells (top panels) or T-bet^+^TNF-α^+^ effector Th1 cells (bottom panels) as determined via FACS analysis after indicated treatments (IL-21, LY 0.025 µM, clotrimazole 10 µM, and 2-DG 5 µM) for 36 h. The bar graph shows the percentage of T-bet^+^IFN-γ^+^ effector Th1 cells (top) or T-bet^+^TNF-α^+^ effector Th1 cells (bottom) (n= 3-4 biological replicates). Data represents mean ± SEM. * p < 0.05, ** p < 0.01, *** p < 0.001 and **** p < 0.0001, using two-tailed Student’s t-test or one-way ANOVA followed by Bonferroni test for multiple comparisons.

Next, we investigated how IL-21 signaling impacts OXPHOS metabolism and T cell immune response. We speculated that IL-21-induced glycolysis and redirection of glucose to the citrate to acetyl-CoA shuttle system would diminish TCA cycle-driven OXPHOS and heighten cytokine production in a manner dependent on GSK3β activity. Consistent with our hypothesis, IL-21 stimulation of human effector Th1 cells diminished glucose-induced basal, ATP-linked and maximal respiration, and spare respiratory capacity (Figures 5C and 5D). Consistent with the data that the IL-21-GSK3β axis (Figures 1K and 1L) triggers OXPHOS to glycolysis metabolic shift, LY2090314 elevated and restored OCR in the absence and presence of IL-21, respectively, compared to vehicle-treated cells (Figures 5C and 5D). Therefore, the IL-21 sustains glycolysis but attenuates OXPHOS in effector Th1 cells in a manner dependent on GSK3β enzymatic activity.

VDAC1 is the main conduit for various respiratory substrates, such as pyruvate, across the outer mitochondrial membrane (OMM) into the inner mitochondrial membrane (IMM). VDAC1-mediated mitochondrial pyruvate influx sustains the TCA cycle and OXPHOS metabolism in cancer cells and cytotoxic CD8^+^ T cells^24, 26, 27^, and GSK3β has been shown to inhibit VDAC1 function via serine-threonine phosphorylation^38^. VDAC1 serine 215 phosphorylation led to VDAC1 dissociation from its activator, mitochondrial hexokinase 1 (HK1), leading to VDAC1 deactivation^39^. Therefore, we speculated that the expression of phosphorylated serine on VDAC1 (pSer-VDAC1) should be evident in effector Th1 cells at steady-state but abolished upon GSK3β genetic ablation. Effector Th1 cells transfected with a CRISPR-Cas9 single guide (sgRNA) that genetically deleted *GSK3B vs*. scrambled sgRNA (Supplementary Figure 5E) were subjected to PLA using monoclonal antibodies directed against VDAC1 and phosphorylated serine to detect and quantify pSer-VDAC1 expression, followed by confocal microscopic imaging of PLA signals within these cells (Figure 5E). Indeed, GSK3β^-/-^ Th1 cells exhibited reduced pSer-VDAC1 expression (Figure 5F, bottom panels) compared to control cells, as evidenced by the loss of red PLA signals (Figure 5F, top panels), indicating that VDAC1 is a substrate of GSK3β. In summary, GSK3β potentially deactivates the VDAC1-mediated mitochondrial conductance of glucose-derived pyruvate via VDAC1 phosphorylation, which may further activate glycolysis needed to sustain histone H3 acetylation and inflammatory cytokine production by effector Th1 cells.

HK1 binding to and activation of VDAC1 at the OMM promotes mitochondrial pyruvate entry and OXPHOS^40–43^, and these mitochondrial signaling events can inhibited by detaching HK1 from VDAC1 via GSK3β activation or pharmacological approaches^24, 29^. Given that effector Th1 cells are capable of both OXPHOS and aerobic glycolysis at steady-state (Figures 1B-1E), we speculated that in a manner dependent on HK1 and GSK3β, VDAC1-mediated mitochondrial conductance of pyruvate governs the balance between OXPHOS and aerobic glycolysis. Furthermore, given that glycolysis is critical for effector Th1 cell inflammatory response, as 2-DG blunted IFN-γ and TNF-α expression (Supplementary Figures 4A and 4B), we hypothesized specific disruption of mitochondrial VDAC1 function in Th1 cells would induce glycolysis switch, leading to amplified immune response. Indeed, VDAC1 inhibition via IL-21-induced activation of GSK3β or exposure to clotrimazole (a fungi-derived product that detaches HK1 from mitochondria, thereby deactivating VDAC1^44^) boosted the percentage of IFN-γ^+^ and TNF-α^+^ Th1 cells (Figure 5G). IL-21 and clotrimazole co-treatment expectedly and synergistically elevated the percentage of IFN-γ^+^ and TNF-α^+^ Th1 cells compared to vehicle or IL-21-treated cells (Figures 5H, 5I, and Supplementary Figure 5F). Moreover, the synergistic inflammatory effect of IL-21 and clotrimazole was blunted with GSK3β inhibitor (LY2090314) or HK2 inhibitor (2-DG), as shown by the reduced frequency of IFN-γ^+^ and TNF-α^+^ Th1 cells (Figures 5H, 5I, and Supplementary Figure 5F). Overall, the IL-21-GSK3β axis in Th1 cells inhibits VDAC1 activity, resulting in OXPHOS to glycolysis metabolic shift and enhanced inflammatory response. These findings led us to assess the therapeutic relevance of restraining GSK3β-dependent processes in a T cell-mediated murine colitis model whose pathology depends on the Treg absence and IFN-γ and TNF-α expression.

### Treatment with GSK3β Inhibitor Prevents Pathogenic CD4^+^T Cell-induced Murine Colitis

So far, our *in vitro* studies suggest that MePyr and GSK3β inhibition undermined the metabolic and inflammatory phenotypes of effector Th1 cells. To demonstrate the potential relevance of Th1 and Th17 cell metabolism to human disease, we first examined whether MePyr and GSK3β-dependent in Th1 metabolic gene signature *in vitro* (Figures 2A-2G and) was enriched in inflamed ileal CD4^+^ T cells from human IBD, a disease characterized by chronic inflammation in and beyond the gut. We re-analyzed single-cell RNA sequencing (scRNA-seq) dataset from human Crohn’s disease ileal lesions in which a pathogenic cell-cell landscape consisting of immunoglobulin <u>G</u> plasma cells, inflammatory <u>m</u>ononuclear phagocytes, <u>a</u>ctivated <u>T</u> cells, and <u>s</u>tromal cells (termed the GIMATS^high^ module) was described to predict patient resistance to anti-TNF therapy^22^. Of note, in FOXP3^-^IL2RA^-^ CD4^+^ T cell clusters from inflamed tissues of GIMATS^high^ patients (i.e., non-responders to anti-TNF) *vs*. GIMATS^low^ patients (responders to anti-TNF), we found upregulation of genes involved in glycolysis (such as *ALDOA*, *GAPDH*, *TPI1*, *PGK1*, *ENO1*, and *LDHB*) and interconnected metabolic pathways, such as nucleotide biosynthesis that consume glycolytic intermediates (*GUK1* and *APRT*), lipid metabolism (*PAFAH1B3)*, and glutaminolysis (glutamine transporter encoded by *SLC38A1*) (Figure 6A). Moreover, these metabolic genes (including *TPI1*, *PGK1*, *ENO1*, *LDHB*, *SLC38A1*, *GUK1*, *APRT*, and *CMPK1*) were also enriched in FOXP3^-^ IL2RA^-^CD4^+^ T cells from inflamed Crohn’s disease tissues *vs*. adjacent uninflamed tissues from the same patients (Figure 6B). Interestingly, glycolytic, biosynthetic, and glutamine metabolism have been described to be critical drivers of Th1/Th17 cell pathogenic function^1, 2, 45^. Overall, the effector Th1 cell metabolic gene signature downregulated by MePyr or GSK3β inhibition *in vitro* is amplified in refractory human IBD.

**Figure 6.**
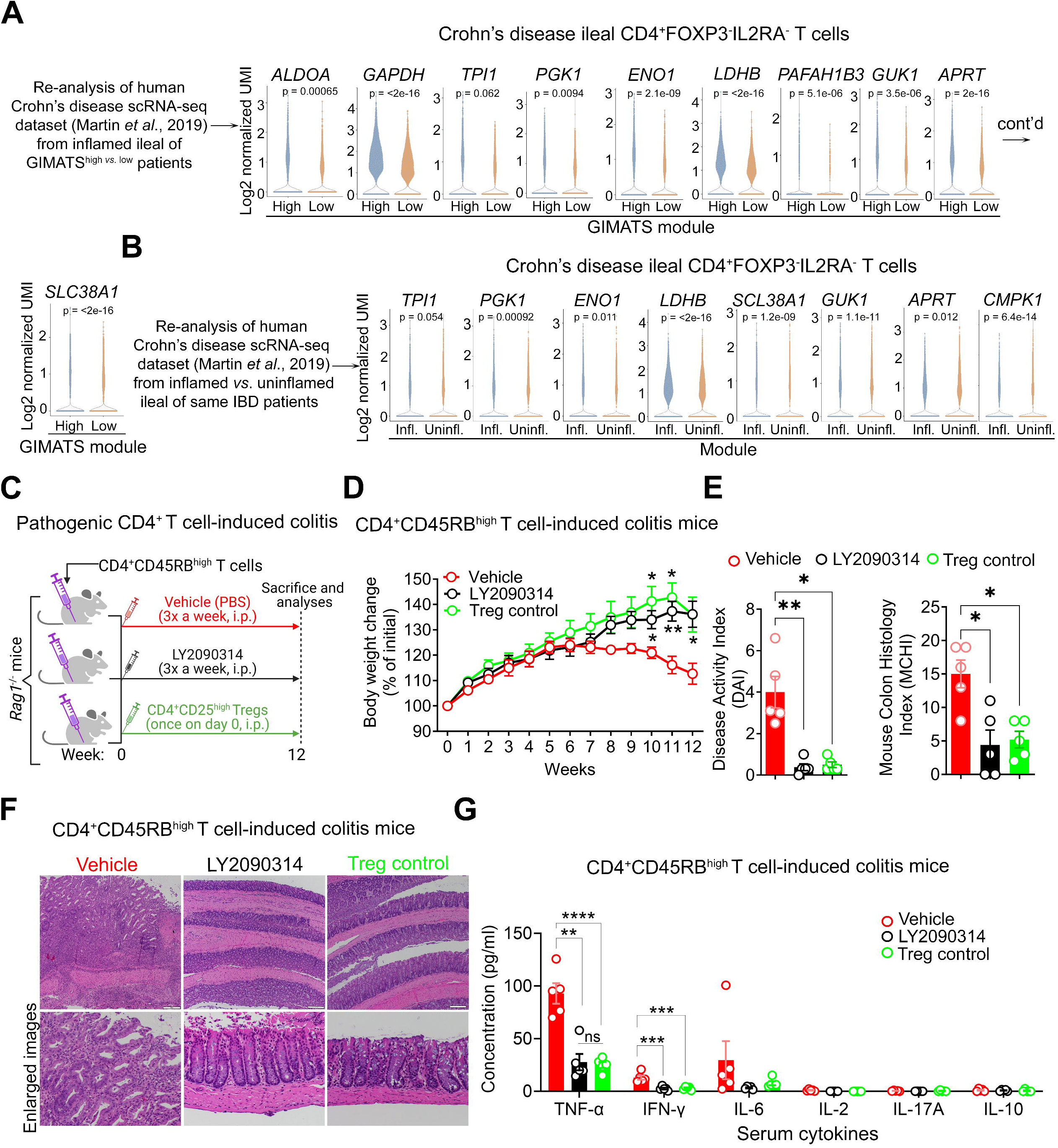
Treatment with GSK3β Inhibitor Prevents Pathogenic CD4^+^T Cell-induced Murine Colitis. (A) Violin plots show the log2 normalized UMI of metabolic genes in inflamed ileal CD4^+^FOXP3^-^IL2RA^-^ T cell (non-Treg) clusters derived from Crohn’s disease of GIMATS (Ig<u>G</u> plasma cells, inflammatory <u>M</u>NP, and <u>a</u>ctivated <u>T</u> and <u>s</u>tromal cells) modules (Wilcoxon test; *P*); n = 5 for GIMATS^high^ patients (non-responders to anti-TNF) *vs*. n = 4 for GIMATS^low^ patients (responders to anti-TNF). (B) Violin plots show the log2 normalized UMI of metabolic genes in CD4^+^FOXP3^-^IL2RA^-^ T cell (non-Treg) clusters derived from inflamed ileal *vs*. the adjacent non-inflamed ileal from the same Crohn’s disease patients (n = 9) (Wilcoxon test; *P*). (C) Schematic illustrates the model of colitis induction in *Rag1*^-/-^ mice via the intraperitoneal (i.p.) injection of CD4^+^CD45RB^high^ T cells followed by disease prevention via the i.p. co-injection of either vehicle, LY2090314 (10 mg/kg), or splenic CD4^+^CD25^++^ T cells (Tregs; 500,000 cells) on day 0. (D) Change in body weight of mice for 12 weeks during colitis progression (n = 5-6 mice per group). (E) DAI (left) and MCHI (right) of treated colitis mice. MCHI was assessed by a blinded pathologist. (F) Hematoxylin and eosin (H&E) staining of colon sections of treated colitis mice on week 12; scale bar, 50 µm. (G) Serum cytokine expression analysis of treated colitis mice on week 12. Data represents mean ± SEM. * p < 0.05, ** p < 0.01, and **** p < 0.0001, using two-way ANOVA followed by Tukey for multiple comparisons, non-parametric Kruskal-Wallis test followed by Dunn’s multiple comparisons (E).

While the etiology of human IBD is complex, the most accredited hypothesis is that this is the result of an inappropriate and exaggerated immune response by T cells^46^. We hypothesized that systemic administration of GSK3β inhibitor would prevent CD4^+^CD45RB^high^ T cell-induced murine colitis. CD4^+^CD45RB^high^ T cells can differentiate into effector Th1/Th17 cells in immunodeficient mice. To address this, we adoptively transferred splenic naïve CD4^+^CD45RB^high^ T cells into mice deficient in the recombinase-activating gene-1 (*Rag1^-/-^*) to induce colitis, followed by co-injection with either the vehicle, GSK3β inhibitor (LY2090314; 10 mg/kg), or splenic Tregs as experimental controls (Figure 6C). Colitis mice injected with the vehicle expectedly lost their body weight (Figure 6D), displayed increased disease activity index (DAI) measuring clinical symptoms, and developed colitis (colonic tissue inflammation and damage) based on a blinded histology assessment defined by mouse colitis histology index [MCHI]) (Figures 6E and 6F) with concomitant elevation of serum inflammatory IFN-γ, TNF-α, and IL-6 cytokines (Figure 6G). Of note, similar to treatment with Tregs, LY2090314 improved weight gain and reduced DAI, MCHI and serum inflammatory IFN-γ and TNF-α cytokines, while IL-2, IL-17A, and IL-10 levels were barely detectable across all groups (Figures 6C-6G).

To provide mechanistic insights into how LY2090314 attenuated CD4^+^ T cell-induced colitis, splenocytes and mesenteric lymph nodes (MLNs) were isolated, activated *ex vivo* with anti-CD3 and CD28 antibodies, and subjected to immunophenotyping via FACS analysis. Like colitis mice treated with Tregs, LY2090314-treated colitis mice displayed a reduced frequency of splenic CD4^+^T-bet^+^ T cells compared to vehicle-treated colitis mice (Supplementary Figures 6A and 6B). However, we did not observe significant changes to MLN CD4^+^T-bet^+^ T cell frequencies across the three groups. Overall, these results suggest that GSK3β enzymatic activity contributes to pathogenic CD4^+^ T cell-induced murine colitis *in vivo*.

## Discussion

This study explored the metabolic mechanisms governing the persistent activation of effector CD4^+^ T cell function. First, effector Th1 cells exhibited elevated glycolysis but low mitochondrial respiration. Enforced mitochondrial metabolism via methyl pyruvate supplementation deactivated GSK3β, resulting in reduced glycolytic function, histone H3 acetylation mark on cytokine promoter region and inflammatory cytokine expression. Second, the metabolic and anti-inflammatory effects of methyl pyruvate on Th1 cells were recapitulated by GSK3β inhibition, and acetate supplementation reversed the anti-inflammatory effects of methyl pyruvate and GSK3β inhibitor. Third, IL-21 reinforced GSK3β, resulting in OXPHOS to glycolysis metabolic shift and an exaggerated Th1 cell inflammatory response in a manner reversible by GSK3β inhibition. Refractory Crohn’s disease is associated with an enrichment of GSK3β-dependent CD4^+^FOXP3^-^IL2RA^-^ T cell glycolytic gene signature. GSK3β inhibition with a repurposed pharmacologic inhibitor (LY2090314) prevented the induction and progression of CD4^+^CD45RB^high^ T cell-mediated colitis in mice.

Activated CD4^+^ T cells producing a plethora of inflammatory cytokines, such as IFN-γ, TNF-α, and IL-17, are enriched in IBD patients’ peripheral blood and intestinal mucosa, leading to the notion that inflammatory T cells may promote resistance to monotherapies against TNF, integrin, and the p40 subunit of IL-12 and IL-23^22, 47, 48^. However, studies describing how these pathogenic immune cells gain inflammatory function to perpetuate IBD have remained elusive. By leveraging findings from the re-analysis of the inflamed ileal scRNA-seq dataset from Crohn’s disease patients and mechanistic *in vitro* and *in vivo* studies, we identified GSK3β as a molecular target that could be inhibited to restrain Th1/Th17 cell inflammatory function and treat human IBD.

Cellular metabolism, guided by nutrient uptake and cues from the microenvironment, is intricately linked to T cell function. We revealed that effector T cells, particularly Th1 cells, displayed superior glycolytic function but low mitochondrial respiration. Forced mitochondrial metabolism via MePyr supplementation diminished glycolysis-mediated histone H3 acetylation of cytokine promoter, which resulted in impaired IFN-γ and TNF-α expression. Mechanistically, MePyr reduced GSK3β activity, resulting in functional restoration of VDAC1, a GSK3β substrate, and the attenuation of HK2-mediated glycolysis with concomitant OXPHOS upregulation. Interestingly, IL-21 enhanced GSK3β activity, possibly via disruption to mitochondria-endoplasmic reticulum interaction as documented in Tregs^29^, which resulted in an intensified Th1 cell inflammatory response. Despite these findings, the molecular intricacies of how GSK3β becomes activated and the mechanism by which MePyr inhibits GSK3β warrant investigation.

The immune system can directly sense the systemic and tissue-specific metabolism to instruct immune alertness required for tissue homeostasis^46^. We observed that sodium acetate supplementation partially restored the inflammatory cytokine-producing capability of methyl pyruvate-treated and LY2090314-treated Th1 cells, supporting the importance of acetyl-CoA in serving as a substrate for histone H3 acetylation needed for Th1 cell immune response. However, it has been reported that acetyl-CoA derived from acetate can also augment glycolysis and effector memory CD8^+^ T cell immune response against systemic acute infection via GAPDH protein acetylation^49^. Excessive glucose uptake by Th1 cells from peripheral blood or gut during the pathogenesis of gastrointestinal disorders, such as IBD, celiac disease, and immune checkpoint inhibitor-induced colitis^50^, may trigger inflammatory immune responses. Furthermore, acetate uptake may potentiate Th1 cell immune response and circumvent the anti-inflammatory effect of IBD monotherapies^13^, leading to therapy resistance and disease perpetuation. Therefore, our study indicates that targeting acetate metabolism in Th1 cells could offer an efficacious therapeutic approach for combating excessive immune response linked to refractory gastrointestinal disorders. Overall, it would be interesting to define how disease-induced alterations to metabolites and cytokines, as reported in the plasma and gut of IBD patients^46^, instruct metabolism and CD4^+^ T cell dysfunction to influence disease severity and biomarker discovery.

Few studies have implicated GSK3β in dextran sodium sulfate and trinitrobenzene sulphonic acid-induced acute murine colitis using other GSK3β inhibitors untested in humans^25^. In this study, we repurposed LY2090314, a GSK3β inhibitor deemed safe in a phase 2 study in acute leukemia patients and unexplored experimentally in CD4^+^ T cells and IBD^30^, to treat T cell-induced colitis in mice. Our study revealed that methyl pyruvate or LY2090314-treated effector Th1 cells gained a Treg-like transcriptional program. Future studies will unexplored whether methyl pyruvate or GSK3β inhibition-mediated activation of OXPHOS-related enzymes can reprogram Th1 cells into Tregs *in vitro* or *in vivo*. Overall, GSK3β inhibition with LY2090314 could, therefore, be a viable therapeutic approach for treating human IBD, either alone or in combination with other therapies.

## Supplementary Figures and Legends

**Supplementary Figure 1. Human Effector Th1 Cells Display High Aerobic Glycolysis but Low Mitochondrial Respiration.**

(A) Relative mRNA expression of *GSK3B* in human effector Th1 cells ± MePyr 10 mM for 36 h as determined via RT-qPCR analysis.

Data represents mean ± SEM. ns (not significant), using a two-tailed Student’s t-test.

**Supplementary Figure 2. Methyl Pyruvate Supplementation Induces Effector Th1 Cell Transcriptional Rewiring via GSK3β Inhibition.**

(A) Venn diagram illustrates upregulated (top) and downregulated (bottom) genes within the 514 detected genes classified as ‘MePyr-dependent genes’ and ‘GSK3-dependent genes’ in Metabolism Panel NanoString analysis.

(B) GO biological process pathway analysis of 16 significant, overlapping upregulated genes in MePyr-treated cells *vs*. LY2090314-treated cells (n = 4) from the NanoString Metabolism Panel using ShinyGO 0.82.

(C) KEGG pathway analysis of 16 significant, overlapping upregulated genes in MePyr-treated cells *vs*. LY2090314-treated cells (n = 4) from the NanoString Metabolism Panel using ShinyGO 0.82.

(D) Venn diagram illustrates upregulated (top) and downregulated (bottom) genes within the 558 detected genes classified as ‘MePyr-dependent genes’ and ‘GSK3-dependent genes’ in Autoimmune Panel NanoString analysis.

(E) GO biological process pathway analysis of 25 significant, overlapping upregulated genes in MePyr-treated cells *vs*. LY2090314-treated cells (n = 4) using the NanoString Autoimmune Panel using ShinyGO 0.82.

(F) KEGG pathway analysis of 25 significant, overlapping upregulated genes in MePyr-treated cells *vs*. LY2090314-treated cells (n = 4) using the NanoString Autoimmune Panel using ShinyGO 0.82.

**Supplementary Figure 3. Methyl Pyruvate Induces Metabolic Rewiring of Human Effector Th1 Cells.**

(A) Experimental workflow for human effector Th17 cell supplementation with MePyr 10 mM *vs*. untreated or treatment with LY2090314 0.025 µM *vs*. DMSO for 36 h, followed by Seahorse analysis.

(B and C) Representative ECAR profile of human effector Th17 cells 36 h after supplementation with MePyr 10 mM compared to untreated cells (n = 3 biological replicates) (B). The bar graphs show calculated glycolysis (basal), glycolytic capacity, and glycolytic reserve; mean ± SEM from 10-12 technical replicates (C).

(D and E) Representative ECAR profile of effector Th1 cells 36 h after supplementation with LY2090314 (LY) 0.025 µM compared to DMSO-treated cells (n = 3 biological replicates) (D). The bar graphs show calculated glycolysis (basal), glycolytic capacity, and glycolytic reserve; mean ± SEM from 10-12 technical replicates (E). Data represents mean ± SEM. **** p < 0.0001 using a two-tailed Student’s t-test. ns (not significant).

**Supplementary Figure 4. Methyl Pyruvate Suppresses Inflammatory Cytokine Production by Effector Th1 Cells via Attenuation of Histone Acetylation.**

(A-B) The representative dot plots show the percentage of human T-bet^+^IFN-γ^+^ (top panels) or T-bet^+^TNF-α^+^ effector Th1 cells (bottom panels) as determined via FACS analysis after exposure to DMSO or 2-DG 10 µM for 36 h, and Fluorescence Minus One (FMO) dot plot indicates the controls used for gating and analysis of double-positive T cell populations (A). The bar graph shows the percentage of T-bet^+^IFN-γ^+^ effector Th1 cells (left) or T-bet^+^TNF-α^+^ effector Th1 cells (right) (n = 4 biological replicates).

(C) The representative dot plots show the percentage of human T-bet^+^IFN-γ^+^ (top panels) or T-bet^+^TNF-α^+^ effector Th1 cells (bottom panels) in response to MePyr 10-20 mM for 36 h in the absence or presence of sodium acetate 20 mM as determined by FACS analysis (n = 3-4 biological replicates).

(D and E) The representative dot plots show the percentage of human T-bet^+^TNF-α^+^ effector Th1 cells in response to LY2090314 (LY 0.025-0.05 µM) or 2-DG 10 µM treatment for 36 h in the absence or presence of sodium acetate 20-50 mM as determined by FACS analysis (n = 3-4 biological replicates), and Fluorescence Minus One (FMO) dot plot indicates the controls used for gating and analysis of double-positive T cell populations (D). The bar graphs show the TNF-α mean fluorescent intensity (MFI; protein expression per cell) in human T-bet^+^TNF-α^+^ effector Th1 cells in response to LY 0.025-0.05 µM or 2-DG 10 µM for 36 h in the absence or presence of sodium acetate 20-50 mM as determined by FACS analysis (n = 3-4 biological replicates) (E).

Data represents mean ± SEM. * p < 0.05, ** p < 0.01, *** p < 0.001 and **** p < 0.0001, using two-tailed Student’s t-test or one-way ANOVA followed by Bonferroni test for multiple comparisons.

**Supplementary Figure 5. IL-21-induced GSK3β activation and Metabolic Rewiring sustain inflammatory Response by Th1 Cells.**

(A and B) Representative ECAR profile of murine WT *vs*. IL-21R^-/-^ effector Th1 cells (cells were pooled from n = 5 mice per group) (A). The bar graphs show calculated glycolysis, glycolytic capacity, and glycolytic reserve; mean ± SEM from 10-12 technical replicates (B).

(C and D) Representative ECAR profile of murine WT *vs*. IL-21R^-/-^ iTregs (cells were pooled from n = 5 mice per group) (C). The bar graphs show calculated glycolysis (basal), glycolytic capacity, and glycolytic reserve; mean ± SEM from 10-12 technical replicates (D).

(E) Representative immunoblot shows GSK3β protein expression relative to β-actin in whole cell lysates derived from effector Th1 cells nucleofected with a scrambled sgRNA *vs*. GSK3β sgRNA.

(F) Fluorescence Minus One (FMO) dot plot indicates the FACS gating strategy used for the analysis of double-positive T-bet^+^IFN-γ^+^ effector Th1 cells (top panels) and T-bet^+^TNF-α^+^ effector Th1 cells (bottom panels).

Data represents mean ± SEM. **** p < 0.0001, using two-tailed Student’s t-test. ns (not significant).

**Supplementary Figure 6. Treatment with GSK3β Inhibitor Prevents Pathogenic CD4+T Cell-induced Murine Colitis.**

(A and B) The representative dot plots show the percentage of CD4^+^T-bet^+^ T cells in activated splenocytes and mesenteric lymph nodes (MLNs) isolated from colitis mice (*Rag1*^-/-^ mice injected with CD4^+^CD45RB^high^ T cells) treated with either vehicle, LY2090314 (10 mg/kg), or splenic CD4^+^CD25^+^ T cells (Tregs); n = 5 mice per group (A). The bar graphs show the percentage of murine *ex vivo* activated CD4^+^T-bet^+^ T cells within the spleen (left) or MLN (right) of colitis mice treated with vehicle, LY2090314 (10 mg/kg), or Tregs; n = 5 mice per group (B). Data represents mean ± SEM. ** p < 0.01 and *** p < 0.001, using two-way ANOVA followed by Tukey for multiple comparisons.

## Acknowledgments

We thank the Mayo Clinic Microscopy and Cell Analysis Core, Mayo Clinic Immune Monitoring Core, and the Metabolomics Core for their experimental and technical assistance.

## Abbreviations

GSK3β: glycogen synthase kinase 3 beta
VDAC1: voltage-dependent anion channel 1
MePyr: methyl pyruvate
2-DG: 2-deoxy-Ɒ-glucose
GAPDH: glyceraldehyde 3-phosphate dehydrogenase
IBD: inflammatory bowel disease
IL: interleukin
i.p.: intraperitoneal
mRNA: messenger RNA
H3K27ac: histone H3 lysine 27 acetylation
H3K9ac: histone H3 lysine 9 acetylation
OXPHOS: oxidative phosphorylation
FCCP: carbonyl cyanide-p-trifluoromethoxy phenylhydrazone
TCA: tricarboxylic acid
ECAR: extracellular acidification rate
OCR: oxygen consumption rate
ETC: electron transport chain
PBMC: peripheral blood mononuclear cell
PLA: proximity ligation assay
scRNA-seq: single-cell RNA sequencing
TCR: T cell receptor
CD3: cluster of differentiation 3
CD28: cluster of differentiation 28
Th: T helper
TNF-α: tumor necrosis factor alpha
IFN-γ: interferon-gamma
Th1: Type 1 T helper
Th17: IL-17-producing T helper
iTreg: induced regulatory T cell
WT: wild-type

