## Supplementary Figures for "Metabolic Reprogramming of Pathogenic CD4^+^ T Helper Cells Attenuates Inflammatory Bowel Disease Pathogenesis"

### SUPPLEMENTARY FIGURE 1

**A**

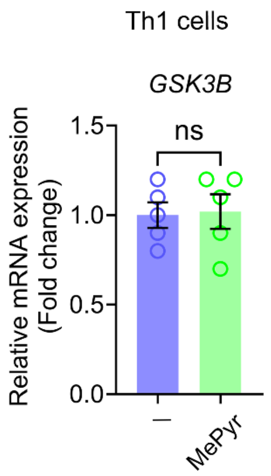

### SUPPLEMENTARY FIGURE 2

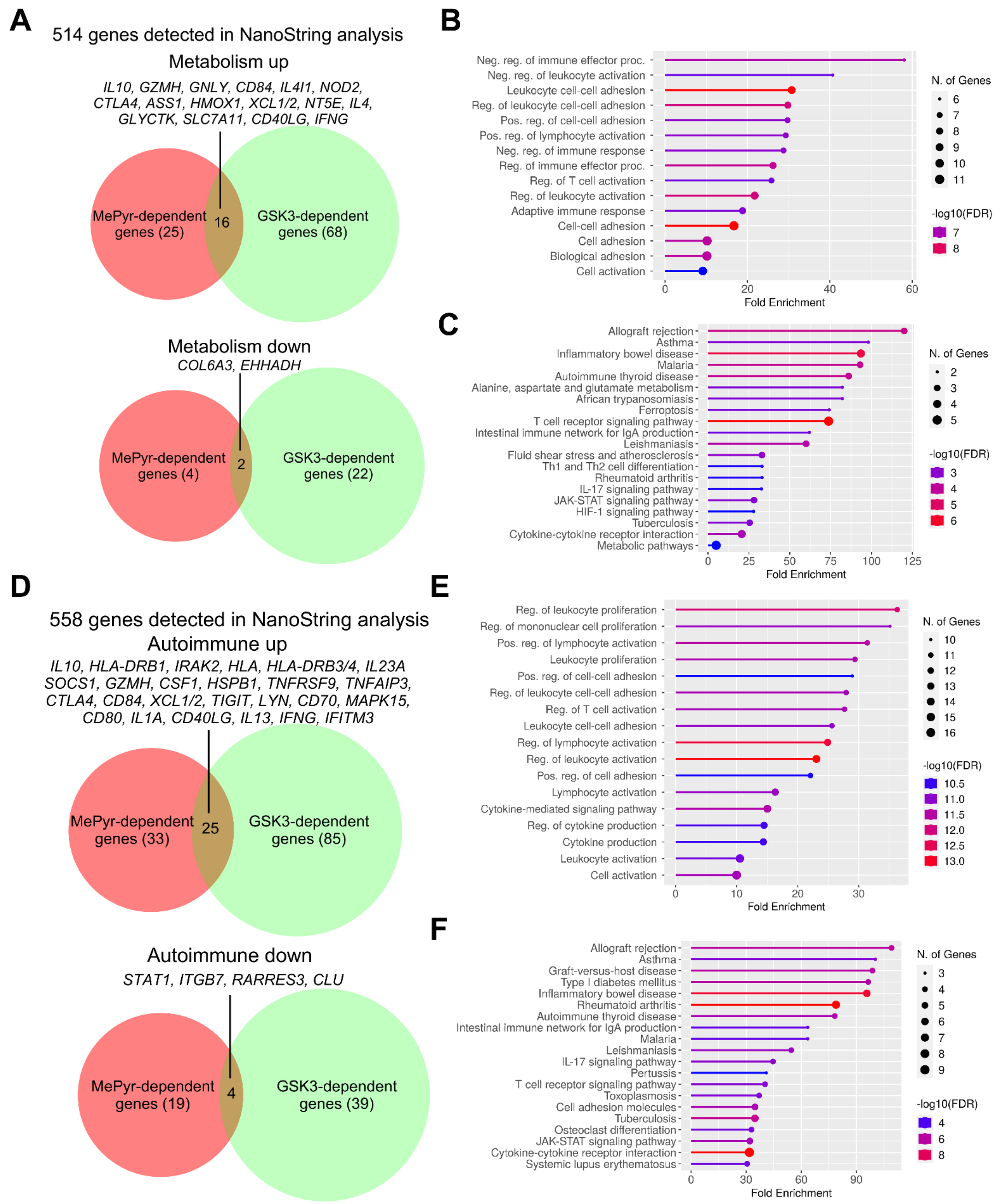

SUPPLEMENTARY FIGURE 3

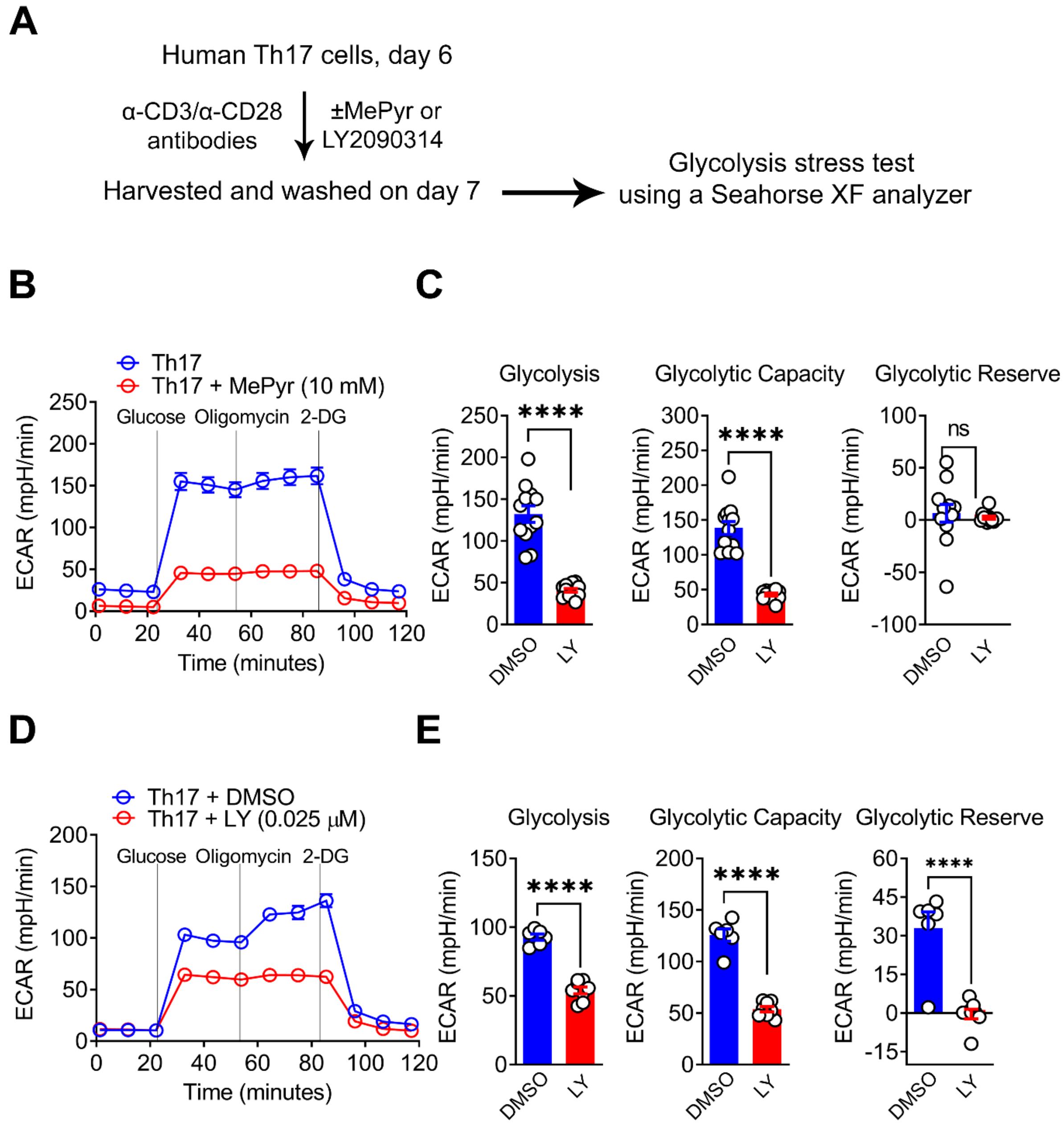

SUPPLEMENTARY FIGURE 4

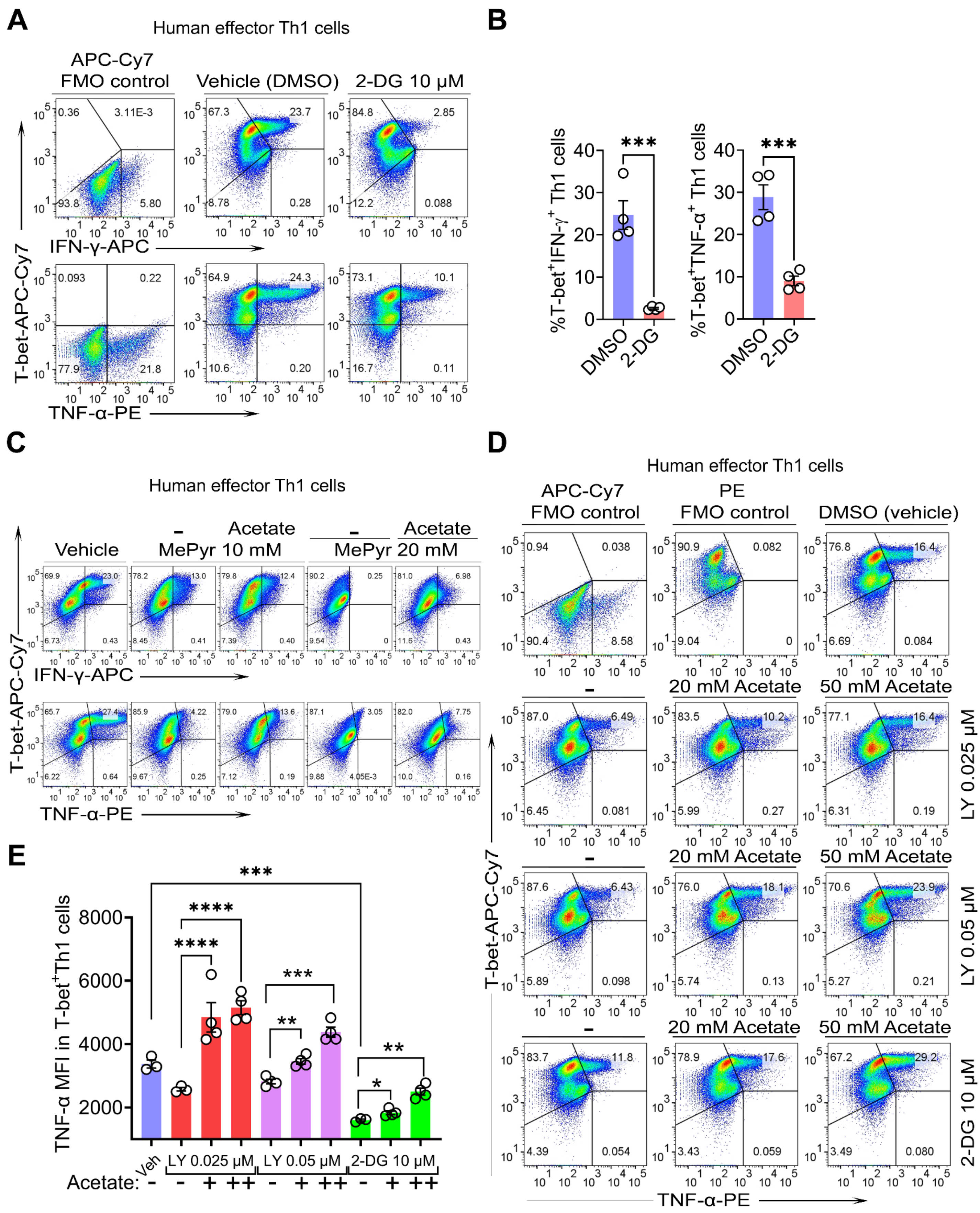

### SUPPLEMENTARY FIGURE 5

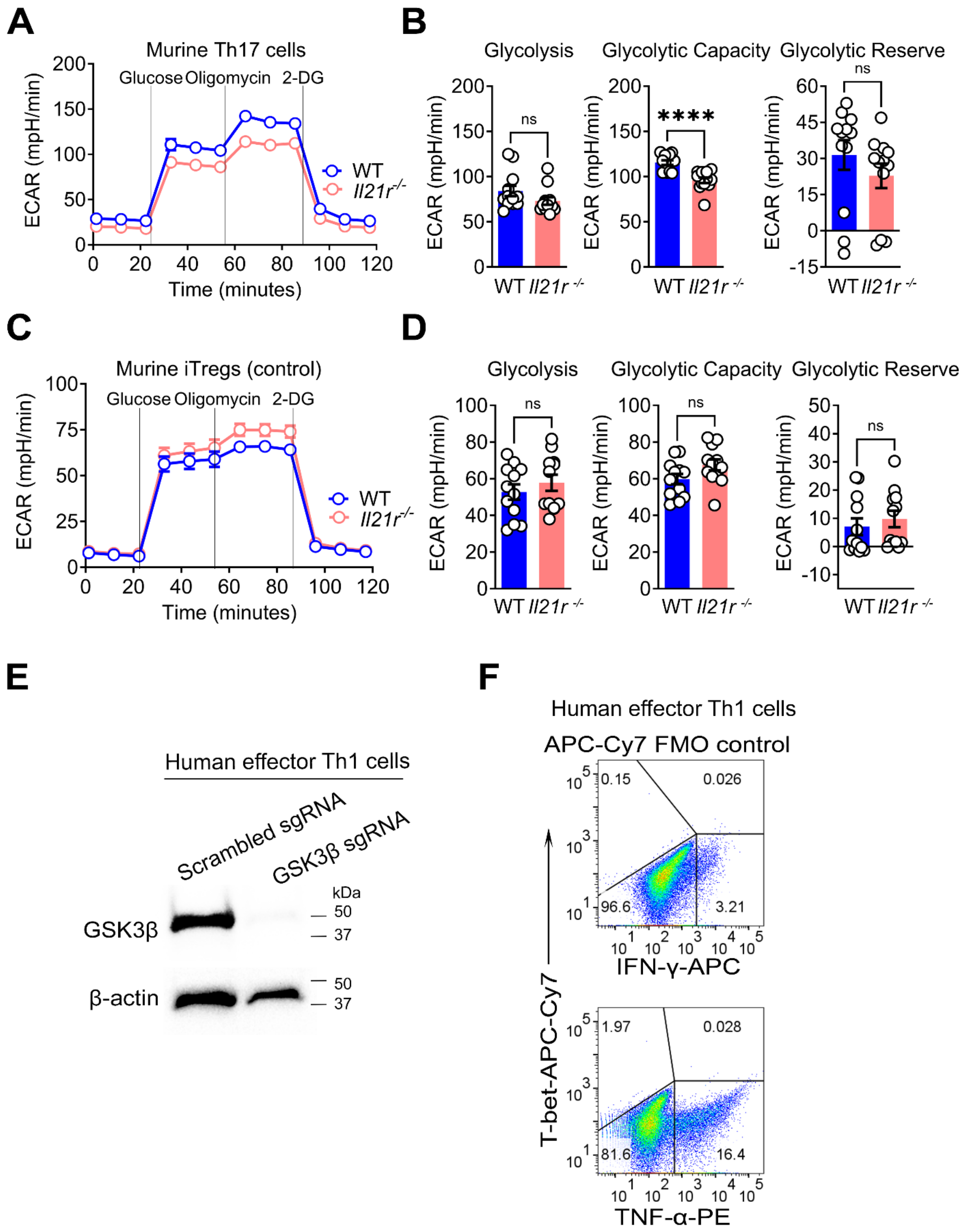

### SUPPLEMENTARY FIGURE 6

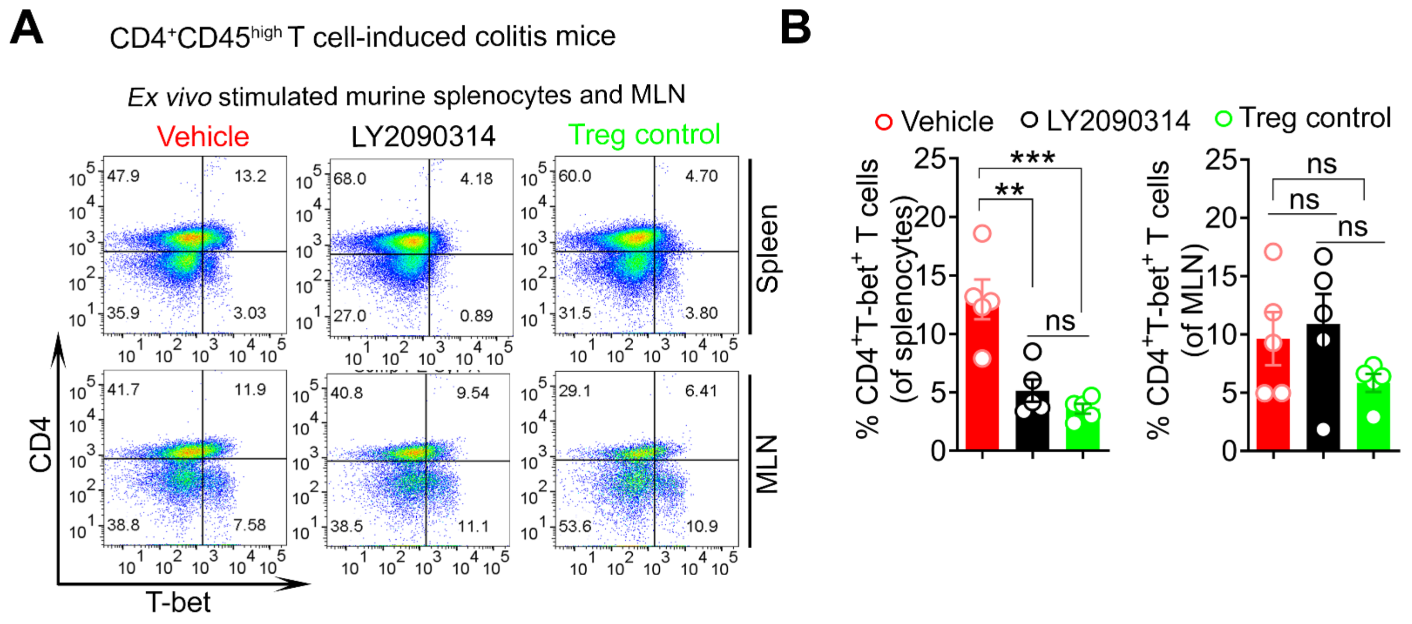
